# Identification and functional characterization of the putative members of the CTDK-1 kinase complex as regulators of growth and development in the genus *Aspergillus*

**DOI:** 10.1101/2023.06.08.544166

**Authors:** Z. Agirrezabala, X. Guruceaga, A. Martin-Vicente, A. Otamendi, A. Fagoaga, J.R. Fortwendel, E.A. Espeso, O. Etxebeste

## Abstract

The genus *Aspergillus* includes industrially, medically and agriculturally important species. All of them, as do fungi in general, disperse to new niches principally by means of asexual spores. Regarding the genetic/molecular control of asexual development, *Aspergillus nidulans* is the main reference. In this species, two pathways control the production of conidiophores, the structures bearing asexual spores (conidia). The Upstream Developmental Activation (UDA) pathway transduces environmental signals, determining whether the Central Developmental Pathway (CDP) and the required morphological changes are induced. The transcriptional regulator BrlA links both pathways as loss-of-function mutations in *flb* (UDA) genes block *brlA* transcription and, consequently, conidiation. However, the aconidial phenotype of specific *flb* mutants is reverted under salt-stress conditions. Previously, we generated a collection of Δ*flbB* mutants unable to conidiate on culture medium supplemented with NaH_2_PO_4_ (0.65M). Here, we identified a Gly347Stop mutation within *flpA* as responsible for the FLIP57 phenotype. The putative cyclin FlpA and the remaining putative components of the C-terminal domain kinase-1 (CTDK-1) complex are necessary for proper germination, growth and developmental patterns in both *A. nidulans* and *A. fumigatus*. Cellular localization and functional interdependencies of the three proteins are also analyzed. Overall, this work links the putative CTDK-1 complex of aspergilli with growth and developmental control.

**One-sentence summary:** Identification of a mutation in *flpA* as inhibitor of conidiation in *A. nidulans* and functional characterization of FlpA, Stk47 and FlpB as putative members of the C-terminal domain kinase complex CTDK-1 in the genus *Aspergillus*.

## Introduction

Aspergilli constitute an important genus of filamentous fungi. Taxonomically, the genus is located within the phylum Ascomycota, subphylum Pezizomycotina and the class of Eurotiomycetes. It is composed of hundreds (∼350) of species [1]. Some of them are pathogens of fruits, seeds, animals and humans. For example, *Aspergillus fumigatus* is the main agent causing invasive aspergillosis [2]. Other aspergilli are used in industry as a source of valuable products [3,4]. *Aspergillus* species can reproduce sexually or asexually [5]. Sexual development has been described in a subset of species of this genus but sequencing of *Aspergillus* genomes has uncovered that the presence of genes coding main regulators of sex is a general trend [1]. Sexual reproduction is based on meiotic cell divisions, generating spores with combinations of the genetic content of the parentals [6,7]. Asexual reproduction is based on mitosis, producing spores with an identical genetic content. This explains why sexual reproduction is mainly directed to the interchange of genetic material while the main aim of asexual development is dispersal.

Spores germinate under favorable environmental conditions. Polar growth and branching of germlings generate mature hyphae, while fusion of hyphae through anastomosis generate the mycelium, the structure specialized in substrate colonization [8]. Depending on the stimulus (O_2_ or CO_2_, light/darkness, nutrient availability/starvation, salt or osmotic stress, presence/absence of specific metabolites) [9] asexual or sexual developmental programs will be activated or repressed. Those spores disperse again to new niches, initiating new life cycles.

One of the main model organisms within the genus *Aspergillus* is *A. nidulans*. Several features make this species a suitable reference organism: fast growth rates at laboratory conditions, the fact that it is not a pathogen, the possibility of carrying out sexual crosses in short periods of time, the amenability for genetic manipulation and the availability of a vast array of standardized cellular and molecular techniques [10]. Together with the Sordariomycete *Neurospora crassa*, *A. nidulans* is the main reference species in the study of developmental programs [3,10,11], since most of the known regulators of sexual and asexual development were identified and characterized for the first time in *A. nidulans* [12]. Asexual spores of *A. nidulans* are known as conidia, since they are produced by localized budding and subsequent constriction from an external sporogenous cell, called the phialide [13]. Phialides and conidia are the last cell-types produced in the asexual developmental cycle, giving rise to multicellular structures called conidiophores. Each *A. nidulans* conidiophore, and the same holds true for *A. fumigatus*, produces thousands of conidia and each colony generates millions of conidiophores on solid culture [9,10,14–16].

In his seminal works, Timberlake estimated that more than a thousand mRNAs increased their concentration after the induction of conidiophore development [17]. To date, the number of identified and functionally characterized proteins with a role in asexual development is far from this estimation. These regulators have been classified into signal transducers, central regulators, repressors and balancers [9,18]. There are complex functional and genetic relationships among these groups of regulators, as well as between them and regulators of other cellular processes such as polar growth, sexual development, secondary metabolism or cell-death. A simplified model describes the activity of two main pathways [19]. UDA (*Upstream Developmental Activation*) pathways are signal transduction pathways involved in the inhibition of polar growth of hyphae and the decision of whether to induce or not, depending on extra- and intracellular stimuli, asexual development. There are at least three UDA subpathways, which are defined by *flbA*, *flbB/D/E* and *flbC*, respectively. FlbB, FlbC and FlbD are transcription factors, and play an important role in the induction of the expression of *brlA*. In fact, *brlA* is the central gene in this model, since its expression is controlled by UDA-s and then, BrlA controls the expression of CDP (*Central Developmental Pathway*) genes, which regulate the formation of most of the cell-types of the conidiophore [9,18,20]. As mentioned above, there are known repressors of *brlA* expression, which act at two levels: repressors that also activate sexual development, and those that repress *brlA* expression once asexual development has been completed [21–25].

Due to their role in the control of *brlA* expression, deletion or loss-of-function mutations in UDA genes inhibit or delay conidiation. Mutants show an aconidial phenotype known as *fluffy* [15]. An example is the Δ*flbB* mutant, in which the CDP pathway is blocked. Nevertheless, the *fluffy* phenotype of the null *flbB* mutant is reverted under specific stress conditions, such as supplementation of the standard solid *Aspergillus* minimal medium (AMM) with high concentrations (above 0.5M) of NaH_2_PO_4_ [26–28]. In a previous work, we took advantage of this phenotype to isolate (through UV mutagenesis) Δ*flbB* mutants unable to conidiate when cultured on AMM supplemented with 0.65M NaH_2_PO_4_ (FLIP, *fluffy* in phosphate, mutants) [28]. Here, mutants FLIP57 and FLIP76 have been characterized. Identification of the mutations causing their phenotypes led to the characterization of AN10640/FlpA (*fluffy* in phosphate A) as a putative cyclin required in the transition from metulae to phialides during conidiophore development. Fluorescence microscopy suggested that its localization is cell cycle-dependent, being located in the nucleoplasm in the interphase but not during mitosis. The same phenotype and subcellular localization were observed for its putative interaction partners AN8190/Stk47, a cyclin-dependent kinase, and AN6312/FlpB, a homolog of *Schizosaccharomyces pombe* Ctk3. Specific dependencies were described among these three proteins for nuclear accumulation and immunodetection patterns. Besides development, germination and growth were also remarkably affected in the null mutants of *flpA*, *stk47* or *flpB*, both in *A. nidulans* and *A. fumigatus*. Overall, results suggest that the activity of the putative CTDK-1 complex in the genus *Aspergillus* is required in multiple cellular processes, participating in the coordination of growth and developmental programs.

## Results

### Sequencing and analysis of FLIP57 and FLIP76 genomes

In a previous work, 80 Δ*flbB* mutant strains unable to conidiate on AMM supplemented with 0.65 M NaH_2_PO_4_ (FLIP mutants) were grouped into seven phenotypic groups [28]. We determined that the mutant phenotype of FLIP166, which belonged to the group of FLIP mutants with a totally aconidial phenotype under phosphate stress conditions, was caused by a mutation in *AN1459/pmtC* [28,29]. Eight additional FLIP mutants were selected (Fig S1), their genomic DNA extracted and *pmtC* sequenced. None of the FLIP mutants analyzed bore a mutation in this gene, showing that their phenotypes were not caused by mutant *pmtC* alleles. Genomic DNA samples of FLIP57 and FLIP76 (Fig 1A) were sequenced and compared to the reference FGSC4 genome, the FLIP166 genome, and *A. nidulans* transcriptomes (see Materials and Methods; see File S1) [24,28,30]. Four and three exonic candidate mutations were identified in FLIP57 and FLIP76, respectively (Fig 1B). However, in FLIP76, an exonic mutation was identified in *AN5717/kapI*, leading to a premature stop codon (R557E+8-Stop; KapI is 1095 amino acids long; File S1; Fig 1B). Since we previously described that the double Δ*kapI*;Δ*flbB* mutant was completely aconidial under stress conditions induced by the addition of 0.5 M NaH_2_PO_4_ [31], we concluded that the FLIP76 phenotype was caused by the additive effect of *flbB* deletion and a loss-of-function mutation in *kapI*. Consequently, we discarded FLIP76 and continued the analysis with FLIP57.

**Figure 1:**
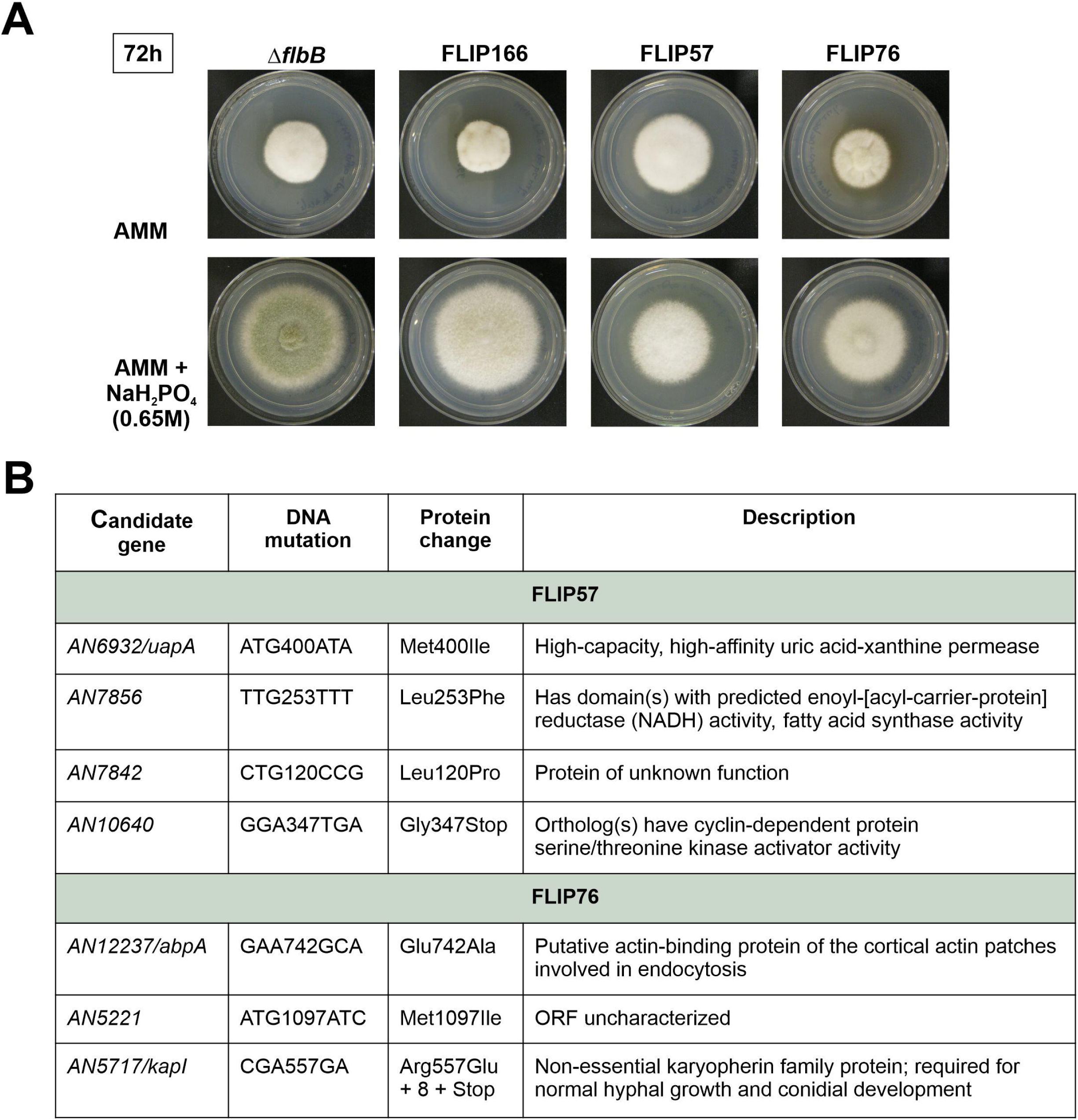
Mutations found in FLIP57 and FLIP76. A) Phenotypes of mutants FLIP57 and FLIP76 after 72 hours of culture at 37 °C in AMM (row 1) or AMM supplemented with 0.65 M NaH_2_PO_4_ (row 2). A parental Δ*flbB* and the mutant FLIP166 [28] were used as controls. B) Mutations found in the genomes of FLIP57 and FLIP76, which could be responsible of the corresponding FLIP phenotypes.

In this second mutant, out of four exonic mutations identified, *AN6932* and *AN7856* (File S1) were discarded because the corresponding null mutant did not show a defect in conidiation [32] or because the gene showed very low expression levels (*AN7856*). *AN7842* was also discarded because, despite the low expression levels, RNA-seq reads suggested that it is not correctly annotated. Thus, the analysis was focused on the mutation found in *AN10640* because: 1) RNA-seq data showed that the annotation of exons and introns was correct, 2) it showed moderate expression levels (see below) and 3) the gene is predicted to encode a cyclin. That would link the control of asexual development with the control of the cell-cycle, transcription and/or additional cellular processes.

### A point mutation corresponding to a Gly347Stop truncation in AN10640/FlpA caused the FLIP57 phenotype

*AN10640* is located in chromosome V (coordinates 1,323,461-1,325,516, coding strand) and is composed of five exons and four introns. The G-T mutation of FLIP57 is located in exon five and modifies the codon in the position 347 (GGA, Gly) by a stop signal (TGA; Gly347Stop) (Fig 2A). Since the corresponding protein is predicted to be 392 amino acids long, the FLIP57 mutation would cause the truncation of the last 45 amino acids (Fig 2A). RNA-seq data showed that, in the culture conditions and genetic backgrounds analyzed by others in *A. nidulans* (reviewed in [12]), the position and extension of introns of *AN10640* matched the FungiDB annotations. Furthermore, RNA-seq data showed moderate expression levels in all conditions/backgrounds analyzed, with no remarkable induction/inhibition in any of the samples (Fig 2B). The Interpro server predicted the presence of a Cyclin-like domain (IPR013763; amino acids 59-263; Fig 2A), which is described to be found in cyclins, but also in transcription factor IIB (TFIIB) and in the retinoblastoma tumor suppressor [33–35]. The FungiDB database described that orthologs of AN10640 have roles in positive regulation of septation initiation signaling, regulation of phosphorylation of RNA polymerase II C- terminal domain and trimeric positive transcription elongation factor complex b localization. The FLIP57 mutation is located outside of this putative Cyclin-like domain.

**Figure 2:**
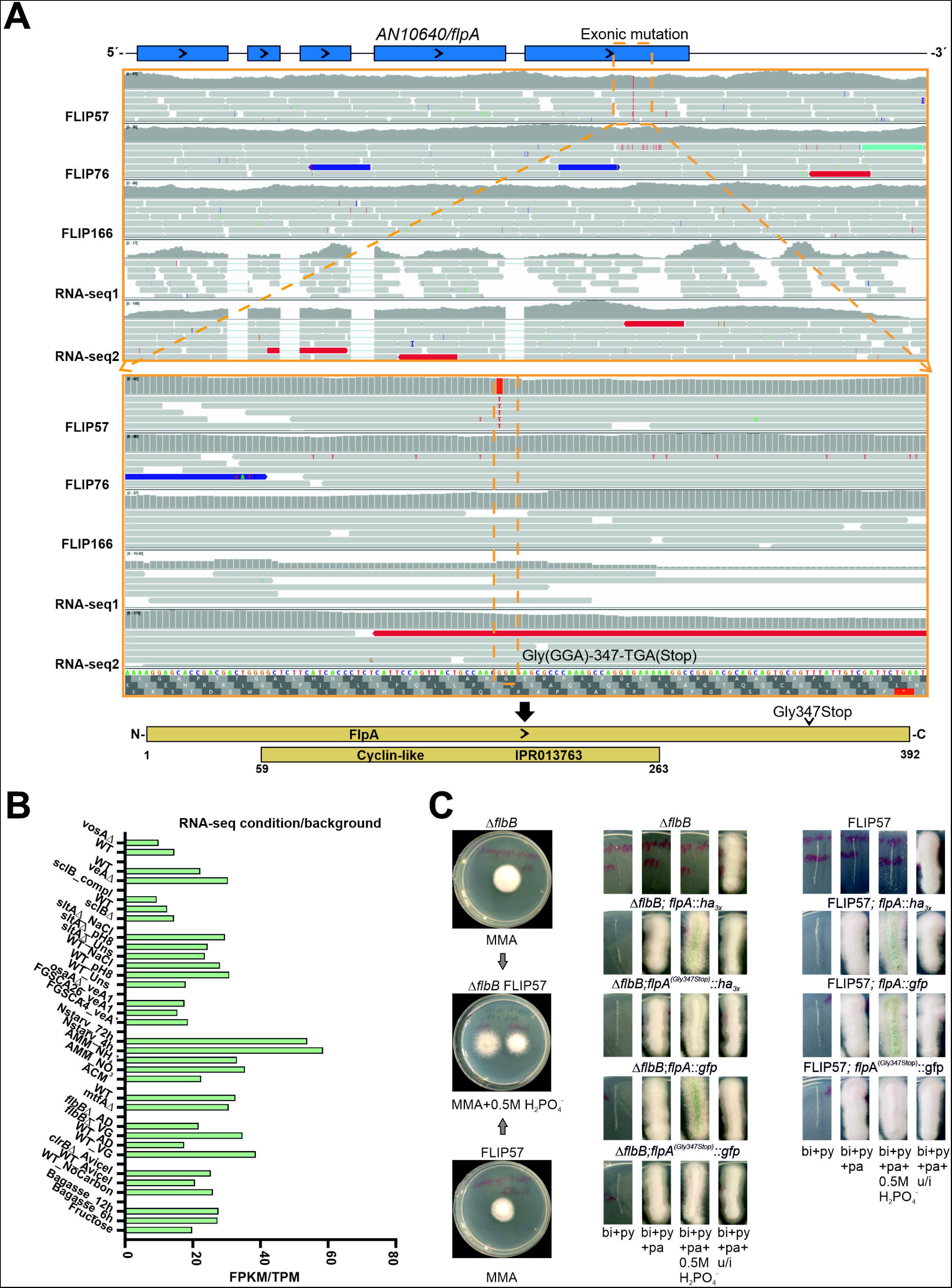
A mutation within *AN10640/flpA* causes the FLIP57 phenotype. A) Annotation of *AN10640/flpA* and analysis of the mutation causing the FLIP57 phenotype. The software IGV was used to visualize the genomes of FLIP57, FLIP76 and FLIP166 and the transcriptomes of a null *flbB* strain (RNA-seq1) and a null *sltA* strain (RNA-seq2) compared to the reference FGSCA4 genome. FLIP57 bore a mutation in the last exon of *AN10640*, which led to the substitution of the codon for Gly347 by a stop codon. This mutation is predicted to truncate the protein and cause the loss of the last 45 amino acids. The identified mutation is outside the cyclin-like domain (IPR013763; amino acids 59-263). B) FPKM or TPM values for *AN10640/flpA* in the multiple RNA-seq experiments available for *A. nidulans* (see references within [12]). C) Confirmation of the Gly347Stop truncation as the mutation causing the FLIP57 phenotype. Left: Phenotypes of FLIP57 and its parental Δ*flbB* strain on AMM and AMM supplemented with 0.5 M NaH_2_PO_4_. Middle: Transformants of Δ*flbB* protoplasts with the synthetic DNA constructs *flpA::ha_3x_::pyrG^Afum^*, *flpA^(Gly347Stop)^::ha_3x_::pyrG^Afum^*, *flpA::gfp::pyrG^Afum^* or *flpA^(Gly347Stop)^::gfp::pyrG^Afum^*. The wild-type *flpA* constructs did not modify the parental phenotype in medium with 0.5 M NaH_2_PO_4_, while the mutant constructs induced a FLIP phenotype. Right: Transformants of FLIP57 protoplasts with the synthetic DNA constructs *flpA::ha_3x_::pyrG^Afum^*, *flpA::gfp::pyrG^Afum^*or *flpA^(Gly347Stop)^::gfp::pyrG^Afum^*. The wild-type *flpA* constructs reverted the parental FLIP57 phenotype in medium with 0.5 M NaH_2_PO_4_, while the mutant construct did not. These results showed that the mutation Gly347Stop within *AN10640/flpA* caused the FLIP57 phenotype.

To confirm that the mutation Gly347Stop in *AN10640* (the gene was named as *flpA*, *fluffy* in phosphate A) was the cause of the FLIP57 phenotype, protoplasts of this strain and those of its parental Δ*flbB* strain were transformed with *flpA::ha_3x_::pyrG^Afum^*, *flpA^(Gly347Stop)^::ha_3x_::pyrG^Afum^*, *flpA::gfp::pyrG^Afum^* or *flpA^(Gly347Stop)^::gfp::pyrG^Afum^*constructs (Fig 2C). The insertion of the mutant constructs in the null *flbB* parental caused an inhibition of conidiation in AMM supplemented with 0.5 M NaH_2_PO_4_ while the wild-type constructs did not alter the Δ*flbB* phenotype. Inversely, the wild-type constructs reverted the FLIP57 phenotype under phosphate stress. No difference was observed between *flpA::ha_3x_* or *flpA::gfp* counterparts, suggesting that the size of the 3’- tag is not detrimental to FlpA activity. On the contrary, the mutant construct *flpA^(Gly347Stop)^::gfp::pyrG^Afum^*did not revert the FLIP57 phenotype under phosphate stress (Fig 2C). These results confirmed the aforementioned hypothesis.

### FlpA and its putative interactors Stk47/AN8190 and FlpB/AN6312 are widely conserved in the kingdom Fungi

Paolillo and colleagues carried out a deep bioinformatics and phylogenetic analysis of *A. nidulans* cyclins [33]. They classified AN10640 within group III cyclins, as a T/K-like cyclin, as was AN4981/PchA. To determine the conservation pattern of FlpA in the fungal kingdom, we carried out BLAST analyses to identify putative ortholog sequences. Table S1 shows the taxonomy of the species corresponding to the FlpA hits analyzed, the length of the hit, score and e values, the query coverage and the result (first sequence) of the confirmatory reverse retrieval of each hit sequence at the FungiDB database. Only hits giving AN10640/FlpA as the first sequence in this confirmatory reverse retrieval were considered. Overall, the values given in Table S1, together with the phylogenetic tree in Fig 3A, show that FlpA is widely conserved in the kingdom Fungi.

**Figure 3:**
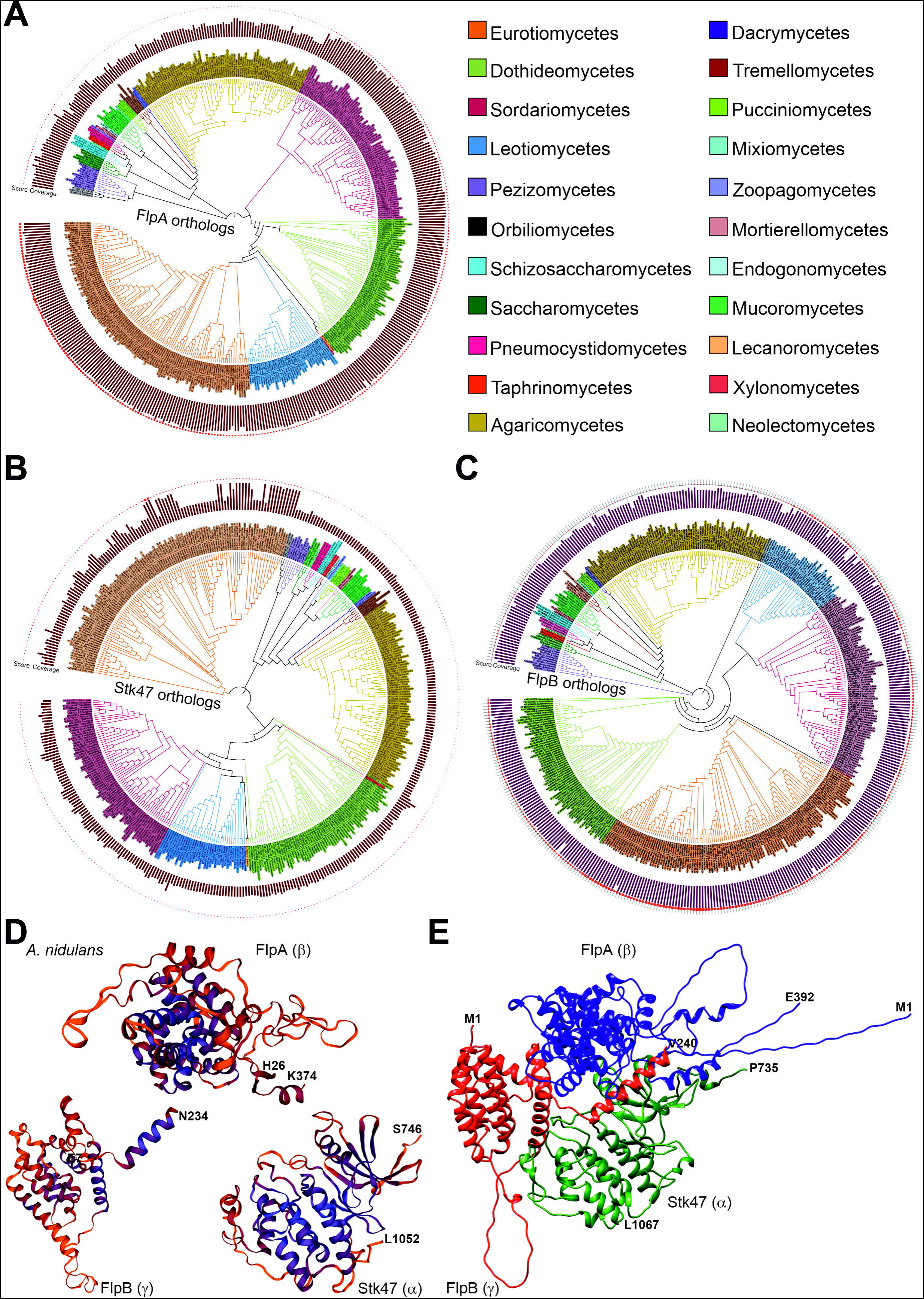
Evolutionary analysis of FlpA/AN10640, Stk47/AN8190 and FlpB/AN6312. Phylogenetic trees corresponding to the putative orthologs of FlpA (A), Stk47 (B) and FlpB (C) included in Table S3. Results show that the conservation of the three subunits of the *S. pombe* CTDK-1 complex is a general pattern in fungi. Ortholog sequences were identified by BLAST analyses. MegaX was used to build the trees (Maximum-likelihood method and the JTT matrix, with 50 replicates in each case) and iTOL to edit them and add coverage (bar graphs in the trees) and score (shape plots) values for each hit compared to *A. nidulans* queries. The color key indicates which fungal class each ortholog belongs to. D) Predicted three-dimensional structures for FlpA, Stk47 and FlpB, modelled by Swiss-Model. The models cover the regions from His26 to Lys374 in FlpA, from Ser746 to His1052 in Stk47 and from Glu7 to Asn234 in FlpB, and are based on PDB structure 7jv7.1, which corresponds to the CTDK-1 complex of *S. pombe* and is shown in Figure S2D [36]. The alignments between FlpA, Stk47 or FlpB and the reference structure can be seen in Fig S2A-S2C. E) Predicted three-dimensional structure for a hypothetic complex formed by FlpA, Stk47 and FlpB, modelled by ColabFold. Full-length sequences of FlpA and FlpB were used, together with an N-terminally truncated form of Stk47 covering residues from Pro735 to Leu1067.

Xie and colleagues have recently determined the structure of the *Schizosaccharomyces pombe* ortholog of FlpA, Ctk2, as part of the trimeric CTDK-1 (C-terminal domain, CTD, kinase I) complex [36]. CTDK-1 acts as the primary RNA polymerase II CTD Ser2 kinase complex. The Swiss-Model website modelled FlpA (from His26 to Lys374; Model 7jv7.1.B was taken as the reference; sequence identity of 18.67%) [36], with higher confidence, as expected, in the cyclin-like domain than in the case of the C-terminus (Fig 3D and Fig S2A). Since it corresponded to a stop signal, Dynamut [37] was unable to predict the hypothetic effect of the FLIP57 mutation in the structure of AN10640.

The remaining two components of the CTDK-1 complex in *S. pombe* are Ctk1, a cyclin-dependent kinase, and Ctk3, the latter being a Ctk1 activator and contributing to the assembly of the complex by interacting with Ctk1 and Ctk2 [36]. The *A. nidulans* ortholog of Ctk1 is AN8190/Stk47 [38] while that of Ctk3 is AN6312/FlpB. Exons and introns of both genes were annotated correctly in the FungiDB database (not shown). Table S1 includes more than 400 fungal species containing FlpA, Stk47 and FlpB (see below) orthologs, representing 22 classes within the phyla Ascomycota, Basidiomycota and Mucoromycota (Fig 3B and 3C). The Swiss-Model website modelled both proteins taking, as in the case of FlpA, PDB structure 7jv7.1 as the reference (subunits α and γ, respectively, for Stk47 and FlpB; much lower coverage in the case of Stk47, from Ser746 to Leu1052; from Glu7- to Asn234 in the case of FlpB) (Fig 3D; Fig S2B-S2C). ColabFold predicted the structure of a putative complex formed by the three *A. nidulans* proteins (full-length versions of FlpA and FlpB were used as queries and the fragment from Pro735 to Leu1067 in the case of Stk47, based in this latter case on the Swiss-Model prediction). The model in Fig 3E shows that FlpB would interact with both FlpA and Stk47, enabling the assembly of the complex, as has been described in *S. pombe* (see the structure in Fig S2D) [36]. Overall, bioinformatics results strongly suggest that CTDK-1 is a widely conserved trimeric complex in the kingdom Fungi while in *Homo sapiens* there is no Ctk3 ortholog [36] (see Discussion).

### Deletion of *flpA* causes a decrease in radial growth and conidial yield, while truncation of the last 45 amino acids of FlpA has negligible effects in a wild-type background

To functionally characterize the role of FlpA in growth and development of *A. nidulans*, a null *flpA* mutant was generated, both in wild-type and Δ*flbB* genetic backgrounds. Wild-type protoplasts were also transformed with wild-type and mutant (Gly347Stop) *flpA::ha_3x_::pyrG^Afum^* constructs, and also a wild-type *flpA::gfp::pyrG^Afum^* construct (see Materials and Methods). Phenotypes of selected strains were analyzed in solid AMM and AMM supplemented with 0.5 M NaH_2_PO_4_ (Fig 4A). First, we observed that, after 72 hours of culture at 37 °C, deletion of *flpA* caused a significant inhibition in radial extension, both in wild-type and null *flbB* backgrounds, and mainly in AMM supplemented with 0.5 M NaH_2_PO_4_ (Fig 4A, 4B and 4C). The average growth rates from 48 to 96 hours of culture were 0.027 ± 0.004 and 0.025 ± 0.003 cm/h for wild-type and Δ*flbB* parentals in medium supplemented with sodium dihydrogen phosphate. The values for the null *flpA* and double null Δ*flpA*;Δ*flbB* strains were 0.012 ± 0.003 and 0.012 ± 0.001 cm/h, respectively, in the same culture medium (n = 3 for each strain; p = 0.006 and 0.004 for each null mutant compared to the parental strain). Furthermore, compared to the null *flpA* mutant (null *flbB* background), the strain that integrated the *flpA^(Gly347Stop)^::ha_3x_::pyrG^Afum^* construct showed only a minor decrease in the growth rate (0.023 ± 0.004 cm/h; n = 3; p = 0.629).

**Figure 4:**
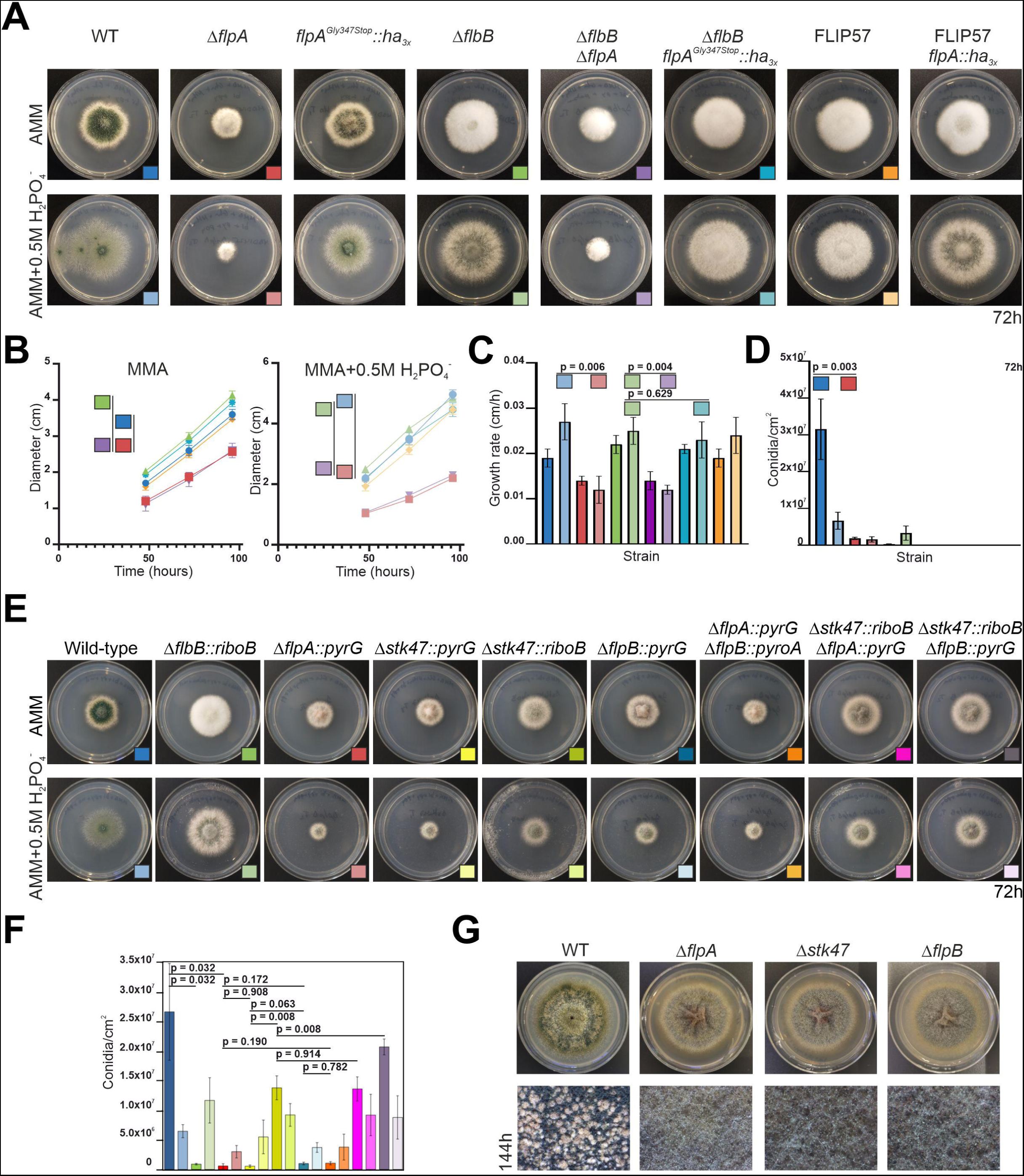
Phenotypic analysis of Δ*flpA*, Δ*stk47* and Δ*flpB* strains in solid media. A) Phenotypes of parental (wild-type, Δ*flbB* and FLIP57) and mutant strains Δ*flpA* (wild-type and Δ*flbB* genetic backgrounds), *flpA^Gly347Stop^::ha_3x_* (same two genetic backgrounds) and *flpA::ha_3x_* (FLIP57 background), after 72 hours of culture at 37 °C in AMM or AMM supplemented with 0.5 M NaH_2_PO_4_. Diameter of Petri plates is 5.5 cm. B) Colony diameters for selected strains (see the color key in panel A) on AMM or AMM supplemented with 0.5 M NaH_2_PO_4_, after 48, 72 and 96 hours of culture at 37 °C. Null *flpA* strains show a clear reduction in colony diameter, mainly under phosphate stress conditions. Each value is the mean of three replicates per strain and time-point, plus SD. C) Growth rates for each of the strains analyzed in panel A. Each value is the average (plus SD) of the growth rates at the three time-points analyzed. The values at each time-point are based on three replicates per strain. D) Conidia production (conidia/cm^2^) for each strain in panel A. Each value is the mean of three replicates, plus SD. Deletion of *flpA* causes a significant inhibition of conidia production. The values of *p* are shown where necessary in panels B, C and D. E) Phenotypes of reference wild-type and Δ*flbB* strains, together to those of single-null *flpA*, *stk47* (generated with *pyrG* or *riboB* markers) and *flpB* mutants, and double-null Δ*flpA::pyrG*;Δ*flpB::pyroA*, Δ*stk47::riboB*;Δ*flpA::pyrG* and Δ*stk47::riboB*;Δ*flpB::pyrG* mutants, after 72 hours of culture at 37 °C in AMM or AMM supplemented with 0.5 M NaH_2_PO_4_. Diameter of Petri plates is 5.5 cm. F) Quantification of conidia production (conidia.cm^-2^) for the strains shown in panel E (see the color key in this panel). *p* values are included for specific comparisons (see main text). G) Inhibition of cleistothecia production in Δ*flpA*, Δ*stk47* and Δ*flpB* strains, compared to their parental wild-type strain, after 144 hours of culture at 37 °C in AMM. Diameter of plates is 5.5 cm. Bottom magnifications correspond to the center of colonies.

The strain that integrated the *flpA^(Gly347Stop)^::ha_3x_::pyrG^Afum^*construct (wild-type background) showed no inhibition of conidia production compared to the reference strain (6.17 x 10^7^ ± 1.45 x 10^7^ and 3.62 x 10^7^ ± 1.55 x 10^7^ conidia/cm^2^, for mutant and wild-type strains, respectively; n = 3 for each strain; mutant not shown in the graphs; p = 0.035) in AMM culture medium. In the null *flpA* strain, however, the inhibition was statistically significant (1.86 x 10^6^ ± 3.23 x 10^5^ conidia/cm^2^ in the null *flpA* strain; n = 3; p = 0.003) (Fig 4A and 4D). The results obtained in the Δ*flbB* background confirmed the trends described for the wild-type background, with the difference that both deletion or truncation of the last 45 codons completely inhibited conidia production, probably due to the additive effect on conidiation caused by the absence of *flbB* [27,31,39]. However, the C-terminus of FlpA could play a minor role in these processes, as observed in the wild-type background.

### Null mutants of *stk47* and *flpB* show the same phenotype as the null *flpA* strain

Since FlpA, Stk47 and FlpB may be the components of the CTDK-1 complex in *A. nidulans*, single-null mutants of *stk47* and *flpB* were also generated and their phenotypes compared to that of the Δ*flpA* strain after 72 hours of culture at 37 °C in solid AMM or AMM supplemented with 0,5 M NaH_2_PO_4_ (Fig 4E and 4F). Different combinations of double-null mutants were also generated by transformation of protoplasts of the single-null mutants with the corresponding deletion cassettes. The three single-null mutants showed a highly similar phenotype. Only in the case of the null *flpB* strain we observed a slight increase in colony diameter and production of conidia. While the reference wild-type strain produced 2.67 x 10^7^ ± 0.82 x 10^7^ conidia/cm^2^, Δ*flpA*, Δ*stk47* and Δ*flpB* strains produced, respectively, 5.94 x 10^5^ ± 3.80 x 10^5^, 5.63 x 10^5^ ± 2.11 x 10^5^ and 1.02 x 10^6^ ± 2.26 x 10^5^ conidia/cm^2^ (p = 0.032 in the comparison between the wild-type and the null *flpA* strains; p = 0.908 and 0.172 when Δ*flpA* was compared to Δ*stk47* and Δ*flpB* strains; p = 0.063 when null *stk47* and null *flpB* strains were compared; n = 3 for each strain). Of note is the phenotypic difference between null *stk47* strains when *A. fumigatus pyrG* or *riboB* genes were used as selection markers (Fig 4E and 4F). Conidia production increased from 5.63 x 10^5^ ± 2.11 x 10^5^ conidia/cm^2^ in the Δ*stk47::pyrG* strain to 1.38 x 10^7^ ± 2.02 x 10^6^ conidia/cm^2^ in the Δ*stk47::riboB* strain (p = 0.008; n = 3 for each strain).

Double-null mutants displayed a highly similar phenotype compared to their corresponding parental strains, and no additive effects in conidia production were observed after deletion of the second gene (Fig 4E and 4F). The double-null Δ*flpA*;Δ*flpB* (generated after transformation of Δ*flpA* protoplasts with the construct for *flpB* deletion) produced 1.07 x 10^6^ ± 2.75 x 10^5^ conidia/cm^2^ (p = 0.190 compared to its parental strain; n = 3 for each strain). Double-null mutants Δ*stk47*;Δ*flpA* and Δ*stk47*;Δ*flpB* (both generated by transformation of protoplast of the Δ*stk47*::*riboB* strain) produced 1.36 x 10^7^ ± 2.04 x 10^6^ and 2.08 x 10^7^ ± 1.34 x 10^6^ conidia/cm^2^ (p = 0.914 and 0.008 when compared to their parental strain).

Finally, the ability to generate cleistothecia by the single-null *flpA*, *stk47* and *flpB* mutants was assessed qualitatively. Fig 4G clearly shows that the mutants do not produce cleistothecia in our culture conditions (7 days of culture at 37 °C in solid AMM). Overall, these results strongly suggest that FlpA, Stk47 and FlpB are necessary for both morphogenesis and sexual or asexual developmental processes. The phenotypic similarities among single-null and double-null mutants also suggest that the three proteins function in the same pathway, in correlation with their hypothetic participation in the formation of the CTDK-1 complex.

### Δ*flpA*, Δ*stk47* and Δ*flpB* strains generate aberrant conidiophores with a deficient metula-to-phialide transition

The null *flbB* mutant fails to induce conidiophore development while deletion of both *flpA* and *flbB* has an additive inhibitory effect on conidia production. Thus, we hypothesized that FlpA could have a role, not at the induction level but, during the generation of the cell types that form conidiophores (foot-cell, stalk, vesicle, metulae, phialides and conidia) [14]. To verify that, we carried out a phenotypic characterization of the Δ*flpA* strain in submerged culture and under nitrogen starvation conditions (see Materials and Methods). As shown before, the wild-type strain produces mature conidiophores after 20 hours of culture at 37 °C under nitrogen starvation conditions (Fig 5A, upper block of images, left) [27,40]. The null *flbB* strain was not able to induce the production of conidiophores (Fig 5A, upper block of images, second column). The Δ*flpA* strain developed conidiophores and conidia under nitrogen starvation. Nevertheless, we repeatedly observed that, in those mutant conidiophores, metulae did not bud correctly into phialides (Fig 5A, upper block of images, right; white arrowheads and magnifications) or that, in specific cases, secondary conidiophores emerged from aberrant primary conidiophores (Fig 5A, lower block of images, column 2). Null *stk47* and *flpB* strains showed the same phenotype as the Δ*flpA* strain (Fig 5A, lower block of images, columns 3 and 4).

**Figure 5:**
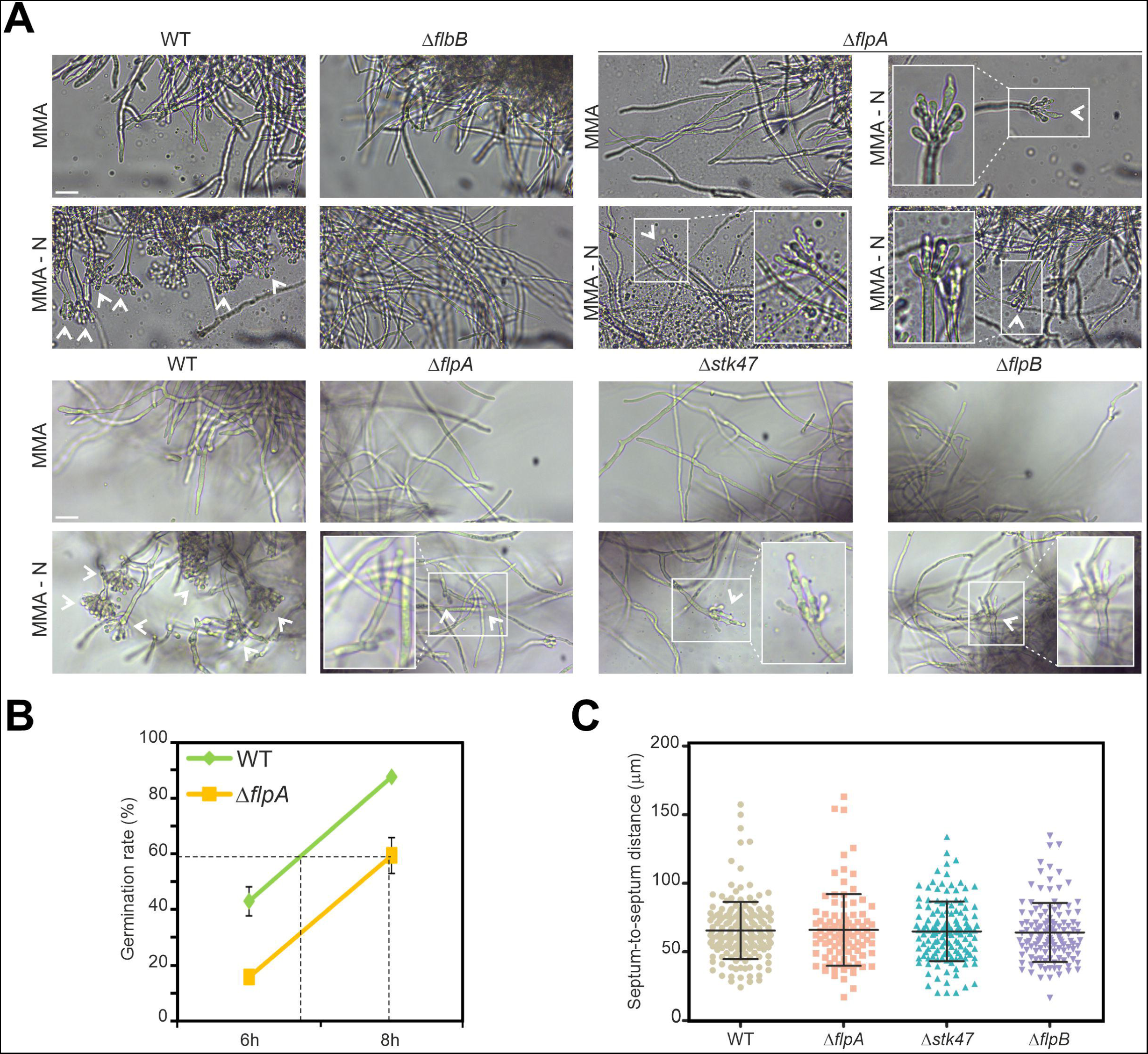
Phenotypic characterization of Δ*flpA*, Δ*stk47* and Δ*flpB* strains in liquid media. A) Up: Phenotype of the Δ*flpA* strain in AMM (18 h + 20 h at 37 °C and 200 rpm; row 1, column 3; see Materials and Methods) and under nitrogen starvation conditions (18 h AMM + 20 h AMM without a nitrogen source; row 2, columns 3 and 4), compared to the parental wild-type strain (column 1) and a Δ*flbB* mutant (column 2). White arrowheads indicate the asexual structures generated by each strain. Those produced by the null *flpA* strain are zoomed to show that most metulae do not bud correctly into two phialides, rendering aberrant conidiophores. Down: Phenotype of the reference wild-type strain and the mutants Δ*flpA*, Δ*stk47* and Δ*flpB* under the same culture conditions as in the upper panel. B) Percentages of wild-type and Δ*flpA* conidia germinated in adequately supplemented AMM (37 °C and 200 rpm), 6 and 8 hours after inoculation (see Materials and Methods). C) Dot-plot showing septum-to-septum distances in null mutants Δ*flpA*, Δ*stk47* and Δ*flpB* compared to the parental wild-type strain. Average and standard deviation values are included. *p* > 0.05 in all comparisons with the wild type.

A hypothetic delay in spore germination was also assessed. Fig 5B shows that spores of the Δ*flpA* mutant are delayed in germination compared to those of the reference wild-type strain. After 8 hours of incubation, 87.6 ± 0.5% of the wild-type spores had germinated, while only 59.3 ± 6.4 % of the null *flpA* spores formed a germ tube (n = 2 replicates per strain), suggesting that there is a delay longer than 1 hour between both strains.

Finally, distribution of septa in hyphae of the null strains of *flpA*, *stk47* and *flpB* was analyzed and compared to that of the parental wild type (Fig 5C). An altered pattern may explain the limited radial growth described previously. The calculated average distance among consecutive septa in wild-type hyphae was 65.6 ± 20.8 µm (n = 182 septa in 41 hyphae). Similar values were measured in the single-null strains, being 66.1 ± 26.1 µm, 65.0 ± 21.7 µm and 64.2 ± 21.5 µm in Δ*flpA*, Δ*flpB* and Δ*stk47*, respectively (n = 97, 141 and 122 septa in 25, 29 and 35 hyphae; p > 0.05 in the three comparisons with the reference strain). Apparently, septa in the null strains were correctly distributed (not shown).

Overall, results explain the reduced radial growth rates and conidia yields described in the previous section and associate these phenotypic traits with a delayed germination and slower radial growth but not an altered distribution of septa, at least in vegetative hyphae.

### FlpA, Stk47 and FlpB localize to nuclei in *A. nidulans* vegetative hyphae

If FlpA, Stk47 and FlpB form the CTDK-1 complex in *A. nidulans* and control the phosphorylation state of the C-terminus of the RNA polymerase II complex, they should localize to nuclei. We generated strains expressing a GFP-tagged version of each protein. When it was driven by the native promoter, FlpA fluorescence could not be detected (Fig S3) and, consequently, a strain expressing a *gpdA ^mini^*-driven [41] FlpA::GFP chimera was generated. FlpA::GFP was detected then in nuclei, probably in a cell cycle-dependent manner, since nuclear accumulation was lost during mitosis (Fig 6A). Stk47::GFP and FlpB::GFP chimeras (driven by their native promoters) were also detected in nuclei of hyphae (Fig 6B and 6C, respectively), confirming our initial hypothesis.

**Figure 6:**
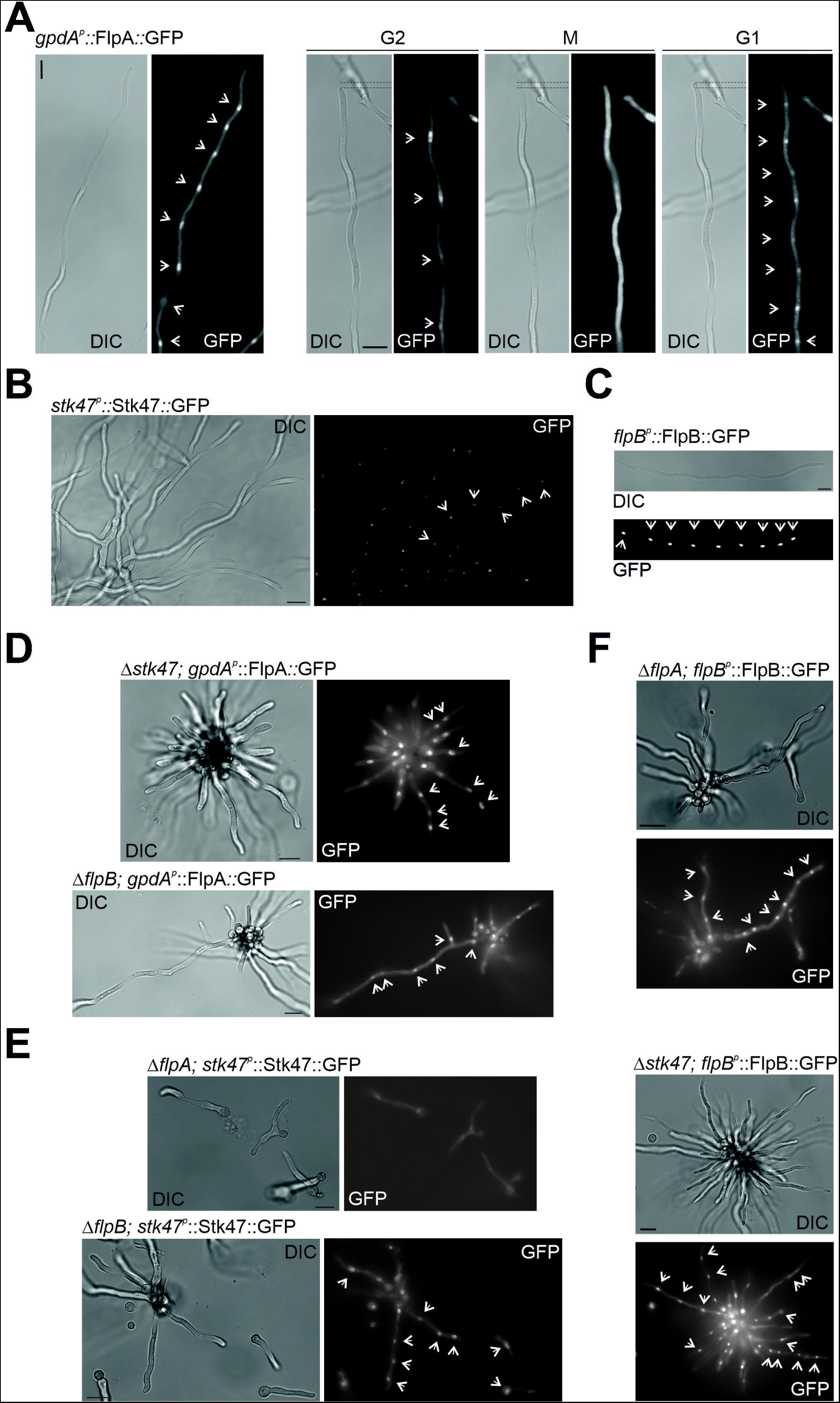
FlpA, Stk47 and FlpB are located in nuclei of vegetative hyphae. Subcellular localization of A) *gpdA^p^*-driven FlpA::GFP (see in Fig S3 the analysis of the same chimera when it is driven by the native promoter), B) Stk47::GFP, and C) FlpB::GFP in vegetative hyphae of a wild-type strain. The nuclear localization of at least FlpA::GFP seems to be cell-cycle dependent, as shown in panel A. D) Subcellular localization of *gpdA^p^*-driven FlpA::GFP in Δ*stk47* and Δ*flpB* backgrounds. E) Subcellular localization of Stk47::GFP in Δ*flpA* and Δ*flpB* backgrounds. F) Subcellular localization of FlpB::GFP in Δ*flpA* and Δ*stk47* backgrounds. Arrowheads highlight the position of nuclei. Scale bars = 10 µm.

Karagiannis and colleagues analyzed the nuclear localization of the *S. pombe* orthologs of FlpA and Stk47, Lsk1 (Ctk1) and Lsc1 (Ctk2) [42]. The authors described that the nuclear localization of Lsc1/Ctk2 was lost in a null *lsk1/ctk1* background, but that in a null *lsc1/ctk2* background, Lsk1 remained at nuclei. We tested any inter-dependence in nuclear localization for FlpA, Stk47 and FlpB with respect to the other potential interactors. With that aim, protoplasts of each single-null mutant were transformed with *gpdA_p_^mini^::flpA::gfp*, *stk47::gfp* or *flpB::gfp* constructs and the subcellular localization of the chimeras was analyzed (Fig 6D, 6E and 6F, respectively). All chimeras accumulated in nuclei, with the exception of Stk47::GFP in a Δ*flpA* background (Fig 6E, upper set of images). These results suggest that the nuclear accumulation of Stk47 depends on FlpA but also that there are additional mechanisms enabling the nuclear import and accumulation of FlpA and FlpB.

### The immunodetection-band pattern of Stk47 is altered in null *flpA*/*B* backgrounds

Next, we carried out immunodetection analyses to determine if the concentration and/or integrity of FlpA, Stk47 or FlpB are dependent on any of the other two activities. FlpA::GFP and FlpB::GFP showed, independently of the genetic background, bands at the expected sizes of 72 and 56 kDa, respectively (Fig 7A). Due to the molecular weight of the Stk47::GFP chimera (155 kDa), we generated strains expressing the Stk47::HA_3x_ chimera (128 kDa) in wild-type, Δ*flpA* and Δ*flpB* genetic backgrounds. We detected two main pairs of bands at different molecular sizes (see red and green squares in Fig 7B). The lightest band in the lower pair of bands (green square) was clearly more intense in Δ*flpA* and Δ*flpB* genetic backgrounds, while the heaviest pair of bands remained unaltered compared to the wild-type background. Despite the detection of these two main pairs of bands, we hypothesized that, considering their high molecular weight, they could hardly correspond to mono- or oligophosphorylation and that they rather corresponded to polyphosphorylation or to a different type of protein modification. Treatment with λ phosphatase (λPP; sodium orthovanadate was used as an inhibitor of phosphatase activity) did not alter the immunodetection pattern of Stk47::HA_3x_ (Fig 7C).

**Figure 7:**
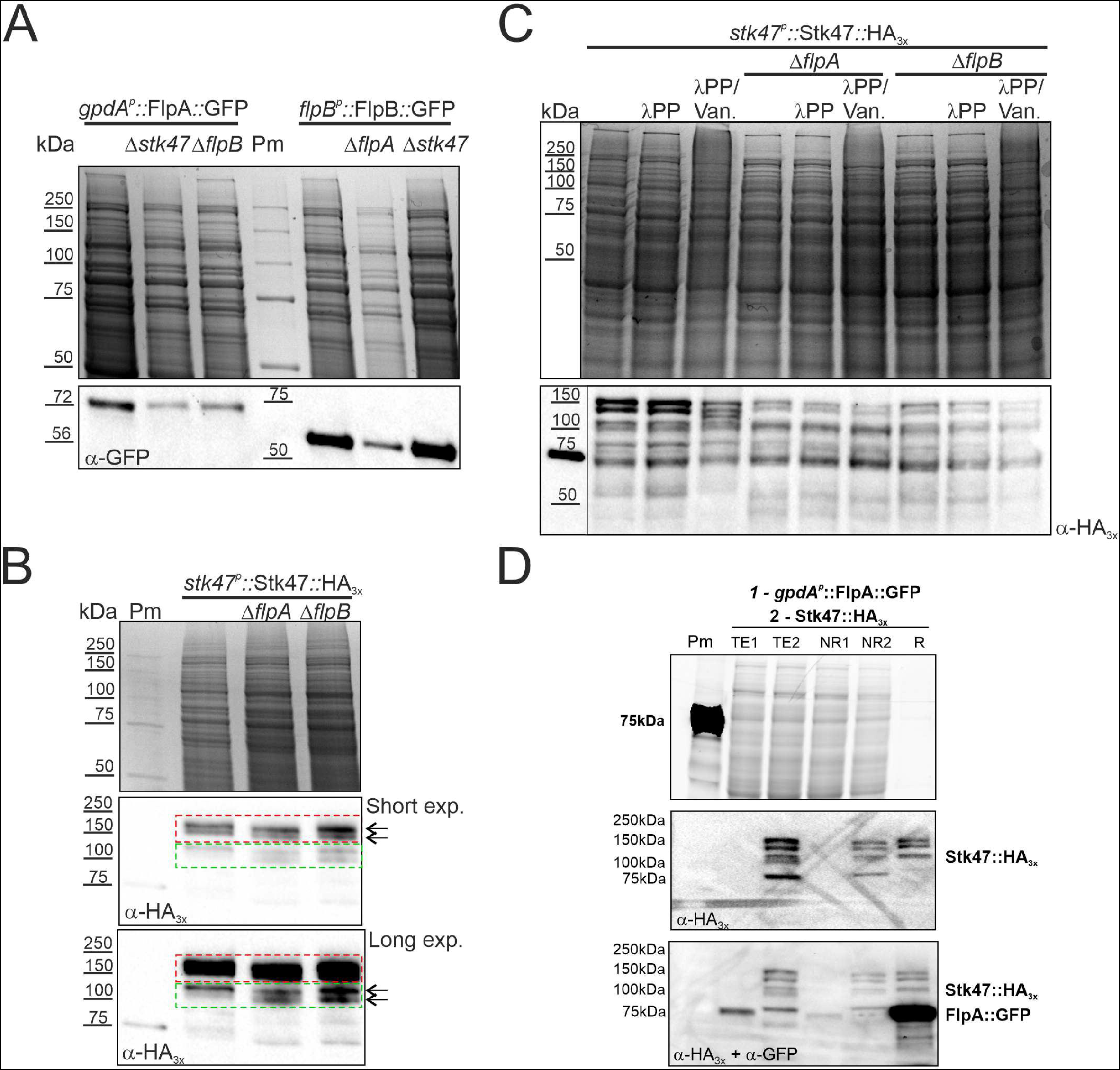
Immunodetection of FlpA, Stk47 and FlpB. A) Immunodetection of FlpA::GFP and FlpB::GFP chimeras, each in the parental wild-type background as well as null backgrounds of the other two genes. B) Immunodetection of Stk47::HA_3x_ in wild-type, Δ*flpA* and Δ*flpB* genetic backgrounds. Stk47::HA_3x_ shows pairs of bands that could correspond to different phosphorylation patterns (red and green squares). Note that within the green square, the lighter band increases its intensity in Δ*flpA* and Δ*flpB* backgrounds. C) Treatment of crude protein extracts of Stk47::HA_3x_ with λ phosphatase (λPP) and λPP plus sodium orthovanadate, the latter as an inhibitor of phosphatase activity. Gels stained with Bio-Safe^TM^ Coomassie (Bio-Rad) are shown as loading controls in panels A-C. D) Pull-down assay using the GFP-Trap resin of Chromotek. Resin samples were sequentially incubated (see Materials and Methods) with 3 mg of a crude protein extract of a strain expressing FlpA::GFP, and after washing of the non-retained fraction, with 3 mg of a second crude protein extract of a strain expressing Stk47::HA_3x_. Strain-Free gels (Bio-Rad) were used for protein electrophoresis. The image in row 1 was obtained as loading control before protein transference to PVDF membranes. TE: Total extract; NR: Non-retained fraction; R: Retained fraction.

Although immunodetection experiments showed an altered Stk47::HA_3x_ band pattern in the single-null mutants of *flpA* and *flpB*, they did not clarify if Stk47 is phosphorylated and if FlpA is the cyclin or one of the cyclins in charge of Stk47 phosphorylation. Thus, we carried out phosphopeptide enrichment analyses to compare the phosphoproteomes of single-null *flpA* and *stk47* strains with that of the wild-type strain (Table S2). Phosphopeptides corresponding to Stk47 were detected in the wild-type strain, but not those corresponding to the large subunit of RNA pol II, Rpb1. The most notably affected function in the phosphoproteome of the Δ*flpA* strain was that of serine/threonine-protein kinases, and this included the absence of phosphorylation of Stk47 (Fig S4). However, results must be taken with care, due to a lack of reproducibility in the second replicate (see Discussion).

To analyze if FlpA and Stk47 form a protein complex, we carried out two types of pull-down assays. First, Stk47::HA_3x_ or (*gpdA_p_*^mini^-driven) FlpA::HA_3x_ chimeras were used as bait for HA-agarose resin (see Materials and Methods). Retained fractions were resolved in SDS-PAGE gels and analyzed through immunodetection. Despite the enrichment in each bait protein (Fig S5), LC-MS/MS analyses of those R fractions did not show significant enrichment in peptides corresponding to putative interactors (FlpA, Stk47 and FlpB) or any other *A. nidulans* cyclin and/or kinases (not shown). Thus, in a second approach, first we incubated the GFP-trap resin with 3 mg of a crude protein extract of a strain expressing *gpdA_p_*^mini^-driven FlpA::GFP (bait; see Materials and Methods). In a second incubation, three mg of a crude protein extract of a strain expressing Stk47::HA_3x_ were added. Fig 7D shows that FlpA::GFP pulls down a fraction of Stk47::HA_3x_.

### Single-null mutants of *A. fumigatus* orthologs of *flpA*, *stk47* and *flpB* show altered growth and developmental patterns

To analyze if FlpA, Stk47 and FlpB activities are also necessary for proper colony growth and developmental patterns in the genus *Aspergillus*, we generated single-null mutants of the three ortholog genes of *A. fumigatus*. Since this species is one of the main animal pathogens in the genus, we also analyzed if these activities are required for growth under stress conditions mimicking the host niche, for altered susceptibilities to antifungal compounds and for virulence in a mouse model of invasive aspergillosis. The three single-null mutants showed a clear growth and developmental phenotype (Fig 8A), with the difference that the radius of Δ*flpB* colonies in medium supplemented with 0.5 M Na_2_HPO_4_ is slightly bigger than those of Δ*flpA* and Δ*stk47* colonies (Fig 8B). In all cases, the decrease in colony *radii* of single-null mutants compared with the reference wild-type strain was statistically significant (p < 0.05 in the three comparisons). We also observed a delay in conidial germination in the three single-null mutants (Fig 8C), which may partially explain the radial growth phenotypes described in Fig 8A and 8B.

**Figure 8:**
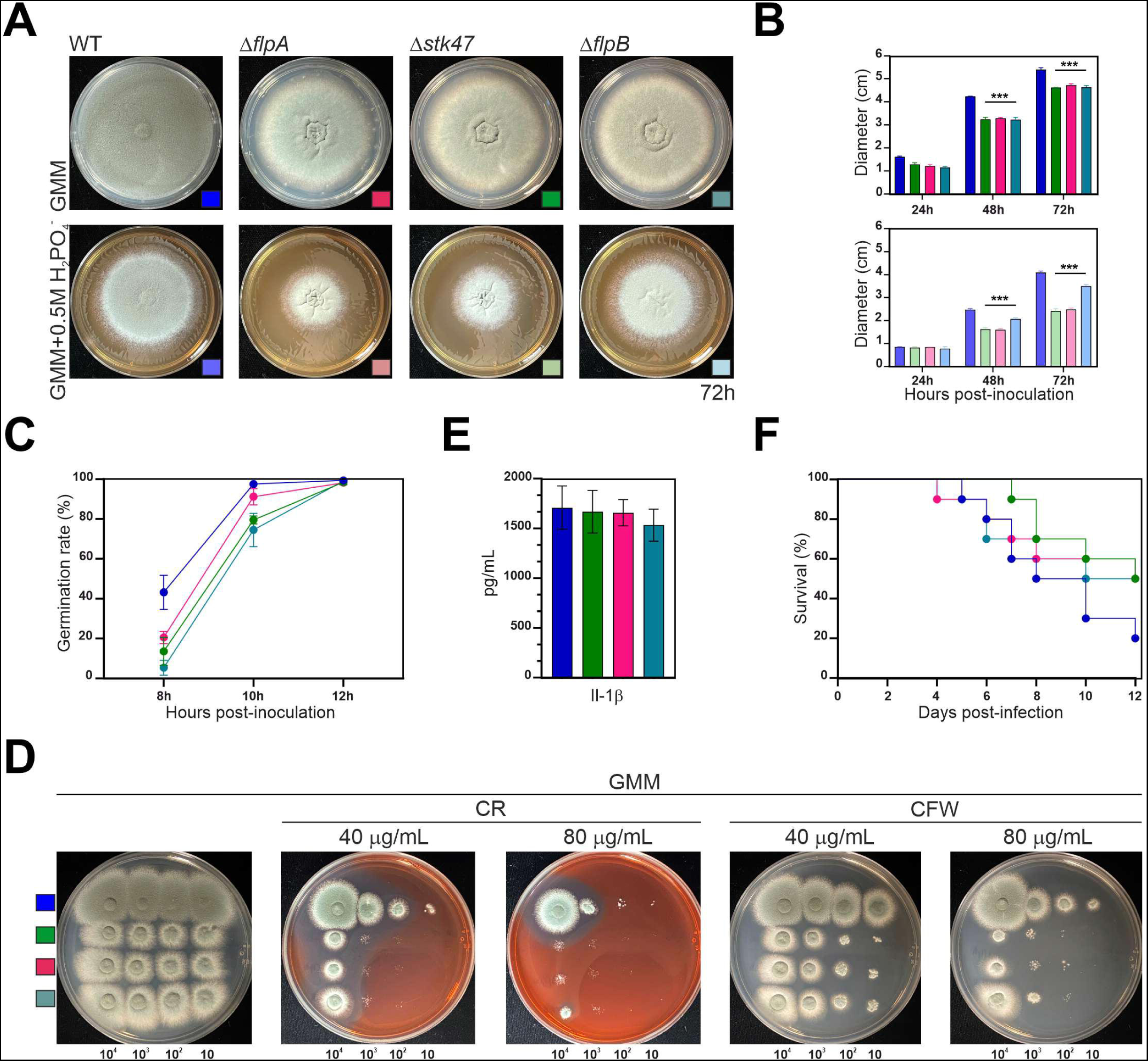
Phenotype of single-null mutants of *flpA*, *stk47* and *flpB* in *A. fumigatus*. A) Phenotypes of wild-type and single-null *flpA*, *stk47* and *flpB* strains in *A. fumigatus* after 72 hours of culture at 37 °C in AMM or AMM supplemented with 0.5 M NaH_2_PO_4_. Diameter of plates is 5.5 cm. B) Diameter of the strains in panel A after 24, 48 and 72 hours of culture under the same conditions as in panel A. Changes in colony diameter of the single-null mutants compared to the reference Δ*akuB-pyrG^+^*strain are statistically significant (***: p < 0.001; n = 3 for each strain). C) Germination rate for each strain 8, 10 or 12 hours after inoculation (n = 3 for each strain at each culture time). D) Phenotypes of the strains after 48 hours of culture on AMM supplemented with 40 or 80 µg.mL^-1^ of Congo-red (CR) or Calcofluor White (CFW). Conidia were inoculated at four different concentrations (10^4^, 10^3^, 10^2^, or 10 conidia). Diameter of plates: 9 cm. E) Quantification of release of interleukin 1β by THP-1 monocytes after co-culturing with conidia (MOI: 2 conidia per THP-1 cell). F) Percent survival of female CD-1 (n = 10 mice per group) to infection by the wild-type or the *A. fumigatus* single-null mutants of *flpA*, *stk47* and *flpB*. Survival was recorded daily and the Kaplan– Meier curves were compared using log rank tests in GraphPad Prism v. 9.2.0.

To test for impacts on growth in host-niche stress, we next examined the ability of the mutants to grow under oxidative, iron limitation, hypoxic and cell wall stresses. Although all strains showed similar growth patterns in response to oxidative stress and under oxygen- or iron-limiting conditions (Fig S6), we found that the Δ*stk47*, Δ*flpA* and Δ*flpB* mutants all exhibited hypersusceptibility to cell wall stress induced by exogenous addition of congo red (CR) or calcofluor white (CFW) (Fig 8D). To see if this cell wall defect would also be evident in response to cell wall-targeting antifungal drugs, we next tested susceptibility of the mutants to the echinocandin caspofungin, which inhibits β-glucan synthesis. Using a strip diffusion assay, we found the wild-type and Δ*flpA* caspofungin MEC was 0.125 µg/mL, and the Δ*flpB* and Δ*stk47* displayed a caspofungin MEC of 0.25 µg/mL (Fig S7). In addition, all strains displayed similar susceptibilities to membrane-targeting antifungals, with a voriconazole MIC range of 0.125 – 0.25 µg/mL and posaconazole MIC of 0.25 µg/mL. Therefore, loss of *stk47*, *flpB* or *flpA* does not impact antifungal susceptibility.

The hypersusceptibility of the three mutant strains to CR and CFW may suggest that the organization of the cell wall, and subsequently PAMP exposure, could be altered by deletion of *flpB*, *flpA*, or *stk47*. Altered PAMP exposure could subsequently lead to changes in host recognition during infection. To check if deletion of any gene alters host recognition, we next challenged THP-1 monocytes with conidia of the fungal strains and assayed inflammasome activation using IL-1β release as an endpoint [43]. However, only a slightly lower release of IL-1β in the single-null mutants compared to the reference wild-type strain was detected and these differences were not statistically significant (p > 0.05 in the three comparisons; Fig 8E). Therefore, we concluded that there are no significant differences in the ability of the mutants to trigger the inflammasome.

Considering the growth, cell wall, and developmental phenotypes, we also analyzed the virulence of the *A. fumigatus* single-null mutants in a chemotherapeutic model of invasive aspergillosis. We observed that, after 12 days of infection, 50% of those animals challenged with the knockouts were alive, compared to only a 20 % of mice infected with the parental strain. However, due to the limited sample size, the mortality rates between strains were not statistically significant (Fig 8F). Taken together with the *in vitro* stress assay data, it is unlikely that *flpB*, *flpA*, or *stk47* play major roles in support of *A. fumigatus* virulence.

## Discussion

Among the different types of kinases, which are differentiated based on the sequence of the kinase domain, cyclin-dependent kinases, or CDKs, are serine/threonine kinases whose activity depends on the regulatory role carried out by a cyclin [44]. Cyclins were named this way because their protein levels fluctuated in a cyclical fashion during the cell cycle and, secondly, because they were defined by the presence of a cyclin box domain (see references within [45]). Cyclins can be divided into two main groups, cell-cycle or canonical cyclins and transcriptional cyclins [45]. The term transcriptional cyclins refers to their role in the regulation of RNA polymerase activity during transcription initiation and elongation. Thus, the role of cyclins goes beyond the control of events directly linked to the cell-cycle. Furthermore, members of both groups of cyclins play cellular functions independently of a partner CDK. Similarly, CDKs also control transcription, metabolism and cell differentiation [46].

Paolillo and coworkers identified, in the genomes of *A. nidulans* and *A. fumigatus*, fifteen genes encoding cyclins that represent the three phylogenetic groups: Group I or cell cycle cyclins (3 in each genome), Group II (7 in each genome) and Group III (5 in each genome), which mainly include transcriptional cyclins [33]. On the other hand, De Souza and Osmani characterized the phenotypes of the single-null mutants of all the *A. nidulans* genes encoding kinases, totaling 128 [38]. The null mutant of *AN8190*/*stk47* was described to show a strong growth defect and sensitivity to stress conditions triggered by sodium chloride. Our groups recently completed the systematic disruption of all putative protein kinase encoding genes in *A. fumigatus* [43]. Interestingly, the *A. fumigatus stk47* disruption mutant displayed only wild-type phenotypes (our unpublished results) which is in contrast to the *A. fumigatus* Δ*stk47* results reported here. However, the previously published work describing these results identified additional kinase disruption mutants that also displayed differential results upon disruption or deletion [43]. These inconsistencies are presumed to be caused by continued transcriptional read through, and subsequent expression of truncated kinase mutants, in at least some of the disruption mutants contained within the *A. fumigatus* library. In the analysis of the set of *fluffy in phosphate* mutants of our *A. nidulans* collection, we identified a Gly347Stop substitution in *AN10640/flpA* as the mutation causing the FLIP57 phenotype. Since FlpA is the ortholog of *S. pombe* cyclin Ctk2/Lsc1, its activity was linked to that of the CDK Stk47 and the hypothetic formation of an *A. nidulans* CTDK-1 complex [36,42]. In humans, CDK12-Cyclin K and CDK13-Cyclin K complexes would be the counterparts of the *S. pombe* Ctk1/Ctk2 complex [36]. Nevertheless, the structure of the *S. pombe* CTDK-1 complex differs from the human counterparts in that it adds a third member, Ctk3, which is not conserved in *Homo sapiens*. Ctk3 enables the formation of the *S. pombe* CTDK-1 complex by interacting with both the cyclin and the kinase. This configuration may be conserved in the kingdom fungi based on: 1) the wide conservation patterns of the three components, 2) the ColabFold prediction, 3) the highly similar phenotypes of the three single-null mutants in *A. nidulans* and *A. fumigatus*, and 4) the interaction pattern described in this work.

The NLStradamus algorithm predicted the presence of a NLS sequence in AnStk47 (1,119 amino acids) but not in AnFlpA (392 amino acids) or AnFlpB (240 amino acids). However, our results showed that, in *A. nidulans*, nuclear accumulation of AnStk47 was dependent on AnFlpA activity while nuclear accumulation of AnFlpA or AnFlpB were not dependent on AnStk47. This behavior is opposite to what was described in *S. pombe*. In fission yeast, Lsk1/Ctk1 was required for nuclear localization of Lsc1/Ctk2, but not inversely [42]. Regarding the phosphorylation pattern of AnStk47, our results are not conclusive. The use of λ phosphatase did not modify the immunodetection-band pattern of AnStk47 and, thus, we were not able to confirm that the Stk47 band pair altered in the Δ*flpA* and Δ*flpB* backgrounds corresponds to a (poly)phosphorylated form of the kinase. Previously, preliminary phosphopeptide-detection analyses by our group (unpublished) detected multiple phosphorylated peptides for AnStk47, suggesting that this kinase may be polyphosphorylated at Ser173, Ser177, Ser221, Ser498, Ser499, Ser505, Ser720, Thr1105, Ser1106, Ser1111 and Ser1112. In addition, the same analysis identified a phosphopeptide phosphorylated both at Ser2 and Ser5 of at least the heptad located at positions 1684-1690 (total length: 1745 aa) of the large subunit of RNA polymerase complex II, AnRpb1. Phosphopeptides for AnFlpA or AnFlpB were not detected. In the phosphopeptide-detection experiments carried out in this work, we were able to detect just a single phosphopeptide corresponding to the hypothetic phosphorylation of Ser720 of AnStk47 (no phosphopeptides for AnRpb1, AnFlpA or AnFlpB were detected). Interestingly, this phosphopeptide was not detected in the null *flpA* strain but, unfortunately, we have not been able to reproduce this result. Dephoure and colleagues described that the probability of observing site-determining ions is hampered by biases in peptide fragmentation that favor breakage at certain points and disfavor it at others [47]. The same authors also stated that the type of protease used in sample processing (trypsin was used in this study; see Materials and Methods) may limit phosphorylation-site identification because not all phosphorylated residues are located in regions that will generate peptides detectable by mass spectrometry upon cleavage by a single protease [47]. Future experiments will have to determine if FlpA is the cyclin or one of the cyclins for Stk47, if both proteins are required for phosphorylation of Rpb1 and if phosphorylation of Stk47 is a prerequisite for its nuclear import and/or retention.

Deletion of *flpA*, *stk47* or *flpB* has pleiotropic effects on colony formation both in *A. nidulans* and *A. fumigatus*. Compared to the reference wild-type strain, 1) germination is delayed, 2) the radius of the colony is decreased, 3) aberrant conidiophores with a misscheduled transition from metulae to phialides are generated, and 4) cleistothecia are not produced. If these three genes encode the components of the CTDK-1 complex, then these phenotypic traits are probably a consequence of an altered phosphorylation pattern of Rpb1. It has been described that phosphorylation of serine 2 of the C-terminal heptad repeats of Rpb1 is not an essential feature of general transcription in fission yeast but rather is critical for certain biological responses, such as sexual development [48]. For example, Ser2 phosphorylation is critical during sexual differentiation of *S. pombe* for the induction of *ste11* transcription [48,49], which encodes an HMG domain mating-type transcription factor. In correlation with these observations, the single-null mutants of *flpA*, *stk47* and *flpB* do not develop cleistothecia, in *A. nidulans*, in the conditions tested. In the case of asexual development, the aberrant conidiophore structures described in this work are probably related to germination and growth defects, rather than 1) misscheduled or incomplete septation processes, as was described in *S. pombe* [42], or 2) a direct role of these three proteins in the control of the expression patterns of UDA or CDP genes. The involvement of genes encoding regulators of germination and hyphal polarity in the formation of conidiophores has been previously described (see for example the cases of *cdc42* and *rac1* orthologs of *A. nidulans* [50]).

The role of transcriptional cyclins and kinases in *A. nidulans* conidiation was previously described by the group of R. Fischer [51,52]. PtkA, a Cdk9 kinase, interacts with cyclins PchA, PclA and PclB, and also with the kinase PipA, modulating its interaction partners and activity to control morphogenesis and development. Cdk9 is supposed to phosphorylate Ser2 residues in the CTD of Rpb1 and enable transcript elongation [53]. Thus, the phosphorylation status of the CTD domain of the large subunit of RNA polymerase II is regulated by multiple cyclin-kinase complexes, which would modify their configuration depending on the morphogenetic/developmental stage and the cell-type. If confirmed in the future for FlpA and Stk47, such a mechanistic flexibility would enable, on the one hand, coordination of multiple pathways and accurate integration of the corresponding signals and, on the other hand, modification of the affinity of RNA polymerase II complex for target promoters.

## Materials and Methods

### DNA sequencing

Genomic DNA of mutants FLIP57 and FLIP76 (Table S3) was extracted following standard procedures and using a phenol:chloroform:isoamyl alcohol mixture (25:24:1 v/v; Pamreac Applichem) [54]. *An1459/pmtC* was amplified from these genomic samples using oligonucleotides PmtC-seq1 and PmtC-GSP2, and sequenced using oligonucleotides PmtC-seq1, PmtC-seq4, PmtC-Up and PmtC-GSP2 (Table S4), to discard that the phenotypes of FLIP57 or FLIP76 were caused by mutations in *pmtC*, as occurred with FLIP166 [28]. After integrity, purity and quantity checks (1.5% agarose electrophoresis and Qubit fluorimeter analyses), DNA samples were sequenced at Stabvida (Caparica, Portugal) using an Illumina Novaseq platform and 150 bp paired-end sequencing reads. The samples generated 16,607,684 (2,507 Mbp) and 16,279,736 sequence reads (2,458 Mbp), respectively, resulting in a theoretical average depth coverage of 83x and 82x, assuming a genome size of 30 Mpb.

### Bioinformatics

Reads were mapped using BWA-MEM [55] at the Galaxy platform (https://usegalaxy.org/), and the SAM files generated were converted to BAM counterparts (SAM-to-BAM). SNPs compared to the reference *A. nidulans* genome (A_nidulans_FGSC_A4_version_s10-m02-r03_chromosomes.fasta) were identified using Snippy. Results for strains FLIP57 and FLIP76 were compared also with those obtained for the mutant FLIP166 [28]. Exonic mutations present in FLIP57 or FLIP76 but absent in the genomes of FLIP166 and the reference genome were selected as candidates. The IGV software (Integrative Genomics Viewer) [56] was used to visualize and analyze DNA- and RNA-seq reads and to compare them with reference genomes.

Gene and protein sequence analyses were carried out in the FungiDB database [57]. The presence of putative functional domains in protein sequences was predicted with InterPro [58], while the NCBI website (https://blast.ncbi.nlm.nih.gov/Blast.cgi) was used to BLAST query sequences. Clustal Omega was used to align protein sequences [59]. The *.msf* file generated was used to visualize the alignments with Genedoc and the *clustal* file generated was imported into MEGAX [60] in order to generate phylogenetic trees, which were edited using iTOL [61]. Structure homology models for FlpA, Stk47 and FlpB were built with Swiss Model [62] and ColabFold [63]. The presence of nuclear localization signals was predicted using NLStradamus and SeqNLS algorithms [64,65]. Finally, GO analyses were carried out at the ShinyGO website (v0.61) [66].

### Strains, oligonucleotides and culture conditions

*Aspergillus nidulans* and *A. fumigatus* strains used in this study are listed in Table S3 while oligonucleotides can be seen in Table S4. Parental strains FLIP57 and FLIP76 were previously generated in a mutagenesis screening of a null *flbB* (BD177) strain [28]. *A. nidulans* strains were cultivated in adequately supplemented liquid or solid AMM, using glucose (2 %) and ammonium tartrate (5 mM) as the sources of carbon and nitrogen, respectively [67,68]. A medium containing 25 g L^−1^ corn steep liquor (Sigma-Aldrich) and sucrose (0.09 M) as the carbon source was used as the fermentation medium (AFM) to culture samples for protein extraction [69]. *A. fumigatus* strains were routinely cultured on AMM agar plates and NaH_2_PO_4_ (0.5 - 0.65 M) was added to AMM to induce salt-stress conditions.

To assess sensitivity of *A. fumigatus* strains to Congo-red (CR) or Calcofluor white (CFW), 10 µL of ten-fold serial dilutions of conidia from each strain (10^4^, 10^3^, 10^2^, 10 conidia) were spotted on Petri Plates filled with AMM supplemented with 40 or 80 µg mL^−1^ of each compound [70]. Plates were incubated for 48 and 72 hours at 37 °C. Colony growth of the mutants was compared to that of the reference strain over the inoculum dilution range.

The ability of the *A. fumigatus* strains to respond to host-stress conditions was evaluated with the following assays: To determine the ability to grow in conditions of iron starvation, 10^3^ conidia were point inoculated in the center of AMM plates or AMM lacking iron, and colony morphologies were observed after 48 hours at 37 °C. To evaluate the growth in low oxygen environments, AMM plates containing 10^3^ conidia of each strain were incubated in a hypoxia chamber at 37 °C for 48 hours. The susceptibility to oxidative stress was evaluated as previously described [71] and growth inhibition halos were measured after 48 hours of incubation. Susceptibility of the strains to posaconazole, voriconazole and caspofungin was determined using gradient diffusion strips (Liofilchem) in RPMI agar plates. Results were read after 24 or 48 hours of growth at 37 °C. All the experiments were performed in biological triplicates.

The procedures developed by Tilburn and colleagues, or Szewczyk and colleagues, were used to obtain and transform protoplasts of *A. nidulans*, and select transformants on selective (lacking uridine and uracil, on the one hand, or pyridoxine, on the other hand) regeneration medium (RMM: AMM supplemented with 1 M sucrose) [72,73]. Protoplasts of *A. fumigatus* strain Δ*akuB*-*pyrG*^+^ were obtained and transformed using the procedure described by Yelton and colleagues [74]. *AfflpA* (*AFUB_007390*; *Afu1g07020*), *Afstk47* (*AFUB_051670*; *Afu5g03160*) and *AfflpB* (*AFUB_027820*; *Afu2g12130*) were deleted using a CRISPR-Cas9 gene editing procedure previously described by us (see below how DNA cassettes were generated) [75]. Selection of transformants was carried out based on hygromycin resistance and correct integration of the DNA constructs was confirmed by diagnostic-PCR using the oligonucleotide pairs shown in Table S4. The control Δ*akuB-pyrG^+^*strain was constructed previously by replacing the mutant *pyrG locus* of the *A. fumigatus KU80*Δ*pyrG* strain with the functional *A. parasiticus pyrG* homologue [70].

For fluorescence microscopy analyses, conidiospores of the strains of interest were incubated for approximately 18 hours at 25 °C in Ibidi µ-Dishes or Falcon multi-well plates containing, respectively, 2 or 1.5 mL of supplemented watch minimal medium (WMM) [76].

To quantify conidia production in *A. nidulans*, asexual spores were point inoculated and cultured for 72 hours at 37 °C. Conidia were collected in Tween 20 (0.02%), diluted (if necessary) and the total amount of conidia determined using a Thoma cell counter. The number of conidia was divided by the area of the colony. Microsoft Excel was used to determine statistically significant differences in conidia production among strains, and to draw the corresponding column bar graphs. Radii of *A. nidulans* colonies and their radial growth rates were determined by measuring and by comparing the diameter of the colonies after 48, 72 and 96 hours of culture in adequately supplemented AMM media.

Conidia of *A. fumigatus* were harvested from 5 day-old cultures grown on AMM. Colony morphologies of *A. fumigatus* mutants were assessed by spotting 5 µL containing 1,000 conidia onto the center of AMM plates (5.5 cm) and incubation for 72 hours at 37 °C. Diameter of colonies were measured after 24, 48 and 72 hours of culture at 37 °C.

The phenotypes of Δ*flpA*, Δ*stk47* and Δ*flpB* mutants of *A. nidulans* under nitrogen starvation conditions were assessed as follows [26,27,40,77]. First, 10^6^ conidia per mL of the mutants and their parental wild-type strain were inoculated in Erlenmeyer flasks filled with 25 mL of adequately supplemented liquid AMM. After 18 hours of culture at 37 °C and 200 rpm, mycelia were filtered using Miracloth paper and inoculated in liquid AMM lacking a nitrogen source. After additional 20 hours of culture, phenotypes were analyzed using an Imaging Source DFK23UP031 digital camera coupled to a Nikon Eclipse E100 microscope.

Hypothetic germination defects of single-null mutants were assessed by inoculating 5 x 10^4^ (*A. nidulans*) conidia per mL in Erlenmeyer flasks filled with adequately supplemented liquid AMM, and by culturing them at 37 °C and 200 rpm. For the evaluation of germination rates in *A. fumigatus*, ten thousand conidia from each strain were inoculated into AMM broth, poured over sterile coverslips and incubated at 37 °C in static conditions. After 6 or 8 (*A. nidulans*), or 8, 10 or 12 (*A. fumigatus*) hours of culture, the germinated conidia were counted from a minimum number of 100 conidia of each strain and time point. At least two replicates were analyzed per strain. The corresponding graphs were drawn with Microsoft Excel or GraphPad Prism v9.2.0.

Distribution of septa in *A. nidulans* hyphae was analyzed using fluorescence brightener 28 (CFW). Briefly, conidia of the strains of interest were inoculated in Falcon multi-well plates containing adequately supplemented WMM and incubated for approximately 18 hours at room temperature. Then, the culture medium was replaced by fresh WMM containing 0.01% CFW. Samples were incubated at room temperature for 5 minutes. Finally, the culture medium was replaced twice (plus additional 5 minutes of incubation) before observation using a Nikon Eclipse Ci manual upright microscope (see below; phase contrast and GFP images were processed). *Septum*-to-*septum* distances (a minimum of 90 for each strain in three biological replicates) were measured using ImageJ (https://imagej.nih.gov/ij/) and the corresponding scatter plot was generated using GraphPad Prism (v5.01).

### Generation of DNA cassettes for transformation

The fusion-PCR technique was used to generate DNA constructs for transformation of *A. nidulans* protoplasts [78]. For 3’-end gene tagging (wild-type or mutant versions), three DNA fragments were amplified and fused. First, 1.5 Kb of the 3’-end of the coding region were amplified using oligonucleotides GSP1 and GSP2 (Table S4). Second, *gfp*, *mCherry* or *ha_3x_* tags plus the selection marker (*pyrG^Afum^* or *pyroA^Afum^* of *A. fumigatus*) were amplified using oligonucleotides GFP1 and GFP2 (2.6 Kb or 1.9 Kb, respectively, when *gfp::pyrG^Afum^*, *gfp::pyroA^Afum^*or *mCherry::pyroA^Afum^*, and *ha_3x_::pyrG^Afum^*were amplified). Third, 1.5 Kb of the 3’-UTR region of the gene (oligonucleotides GSP3 and GSP4). The fusion-PCR reaction was carried out with an aliquot of each fragment and oligonucleotides GSP1 and GSP4.

To generate the constructs for *A. nidulans* gene deletion, the following three fragments were amplified and fused. First, 1.5 Kb of the promoter region of the gene were amplified using oligonucleotides PP1 and PP2. Second, the selection markers *pyrG^Afum^* or *pyroA^Afum^* were amplified with oligonucleotides SMP1 and GFP2. Third, 1.5 Kb of the 3’-UTR region of the gene were amplified with oligonucleotides GSP3 and GSP4. The fusion-PCR reaction was carried out with oligonucleotides PP1 and GSP4. To constitutively express FlpA::GFP or FlpA::HA_3x_ chimeras under the control of the “*mini*” version of the promoter *gpdA^p^* [41], five fragments were amplified and fused (Otamendi *et al.*, 2019b; Otamendi *et al.*, 2019a). First, 1.5 Kb of the promoter using oligonucleotides PP1 and PP2’-ATG. Second, the *gpdA^p^* promoter with oligonucleotides gpdAUp and gpdADw. Third, the coding region of *flpA* was amplified with oligonucleotides geneSP and GSP2. Fourth, the tag plus the selection marker (GFP1 and GFP2). And finally, 1.5 Kb of the 3’-UTR region with oligonucleotides GSP3 and GSP4. The fusion of all those five fragments was done using oligonucleotides PP1 and GSP4. After transformation of *A. nidulans* protoplasts and genomic DNA extraction, correct recombination of the constructs was confirmed by diagnostic PCR using oligonucleotides sPP1 and sGSP4.

For *A. fumigatus* genetic engineering, hygromycin resistance repair templates, containing 40 bp micro-homology regions at both 5’ and 3’ ends of *Afstk47*, *AfflpA* or *AfflpB*, were amplified from plasmid pJMR2 [80] and utilized in a CRISPR/Cas9- mediated transformation to generate complete gene deletions [81].

### Analysis of cytokine release in THP-1 cells

Measurements of IL1-β released by THP-1 cells was carried out following procedures described previously [43,82,83]. THP-1 cells were cultured in RPMI-1640 medium containing HEPES (25 mM) supplemented with heat-inactivated Fetal Bovine Serum (10 %), penicillin and streptomycin (100 U/mL), and 100 μg/mL normocin, and assayed for viability by exclusionary Trypan Blue straining. Plating was carried out at a density of 5 x 10^4^ cells per well (96-well plates) in the same medium but lacking normocin. PMA (phorbol 12-myristate 13-acetate) at a final concentration of 100 nM was added and cells were incubated for 24 hours to differentiate to a macrophage phenotype. Supernatants were discarded and replaced with 180 μL of fresh RPMI (without phenol red) and 20 μL of distilled water containing 5 x 10^5^ *A. fumigatus* conidia (MOI, multiplicity of infection, 10:1) of the single-nulls or the reference strain. Cells were co- cultured for 16 hours at 37 °C and 95% humidity. After incubation time, the supernatants were collected and we measured the IL-1β concentrations using IL-1β ELISA kits (Invitrogen) and following manufacturer instructions.

### Virulence assays in *A. fumigatus*

All animal experiments were carried out according to the protocols approved by the University of Tennessee Institutional Animal Care and Use Committee. Groups of 10, six-week-old female CD-1 mice (Charles Rivers Laboratories) were immunosuppressed with 150 mg/Kg of cyclophosphamide (neutropenic model) on day -3 and subsequently every third day starting again on day +1 [84] and an additional subcutaneous injection of triamcinolone acetonide on day -1. On the day of the infection, mice were slightly anesthetized with 3.5 % isoflurane and inoculated intranasally with 10^6^ freshly harvested conidia contained in 35 µL of pyrogen-free saline solution. Mice were monitored at least twice a day and were given food and water *ad libitum*, and those animals showing severe signs of infection were humanely euthanized by anoxia with CO_2_. Mortality was monitored for 12 days. GraphPad Prism 9.2.0 was used to plot survival data on a Kaplan-Meier curve and to carry out Log-rank (Mantel-Cox) tests.

### Fluorescence microscopy

Subcellular localization of FlpA, Stk47 and FlpB were analyzed using a Zeiss Axio Observer Z1 inverted microscope as previously described [28,85], or a Nikon Eclipse Ci manual upright microscope. The former was equipped with a 63× Plan Apochromat 1.4 oil immersion lens, and filters 38 (excitation at 470 nm and emission at 525 nm) and 43 (excitation at 545 nm and emission at 605 nm). The latter included 40× and 100× (oil) Nikon CFI Plan Fluor lenses, and Semrock filters DAPI-5060C (excitation at 377 nm and emission at 447 nm), GFP-4050B (excitation at 466 nm and emission at 525 nm) and mCherry-C (excitation at 562 nm and emission at 641 nm). ImageJ (https://imagej.nih.gov/ij/) (US. National Institutes of Health, Bethesda, MA, USA) was used to process fluorescence microscopy and phase contrast images.

### Protein Extraction and Immunodetection

For protein extraction, conidia of the strains of interest were collected in tween 20 (4 mL; 0.02 %) and washed twice by centrifugation at 4000 rpm for 10 minutes. By default, inocula (10^6^ conidia mL^-1^) were cultured in AFM (or AMM for phosphopeptide enrichment analyses) for 15 h at 37 °C and 200 rpm [86]. Mycelia were filtered, frozen, lyophilized, homogenized and resuspended in 1 mL per sample of A-40 buffer (25 mM HEPES pH = 7.5, 50 mM KCl, 5 mM MgCl_2_, 0.1 M EDTA, 10 % glycerol and 0.5 mM DTT, supplemented with a protease inhibitor cocktail from Roche) or *Trap* buffer (10 mM HEPES pH = 7.5, 1 mM EDTA, 0.1 % NP-40, 150 mM NaCl, 0.5 mM DTT, plus a protease inhibitor cocktail from Roche; used in pull-down experiments with the GFP-Trap resin of Proteintech). After incubation at 4 °C during 90 minutes with gentle end-over-end mixing, samples were centrifuged at 4 °C and 13,000 rpm for 30 minutes. Supernatants were transferred to new 1.5 mL tubes and protein concentrations were determined by the Bradford assay. Two hundred micrograms of protein were precipitated per sample with trichloroacetic acid (TCA) and purified with ethanol/eter 1:1 and 1:3 mixes, respectively. Finally, precipitates were resuspended in 80 µl of SDS-PAGE loading buffer (62.5 mM Tris–HCl pH = 6.8, 2% SDS, 5% β-mercaptoethanol, 6 M urea, and 0.05 % bromophenol blue) and the integrity of the samples was assessed by polyacrylamide gel electrophoresis.

Alternatively, the procedure described by Hervás-Aguilar and Peñalva was followed [87]. Mycelial samples (approximately 6 mg of lyophilized powder) were lysed using 1 mL of alkaline lysis buffer (0.2 M NaOH and 0.2 % β-mercaptoethanol), and proteins were precipitated by adding 7.5 % TCA. Samples were centrifuged and the supernatants discarded, resuspending the pellets in 100 μL Tris-Base (1 M) and 200 μL of SDS-PAGE loading buffer. Aliquots of 10 µL were then loaded and separated in 10% polyacrylamide gels.

Samples of a hundred micrograms of protein from the crude extracts of the strains of interest were incubated with Lambda phosphatase (λPP; 400 u per reaction; New England Biolabs) for 20 minutes at 30 °C [88]. Sodium orthovanadate (10 mM) was used as a λPP inhibitor when necessary. After incubation, proteins were precipitated as described above and resuspended in 50 µL of SDS-PAGE loading buffer. The integrity of samples was checked by polyacrylamide gel electrophoresis.

For immunodetection, protein samples were separated in Any kD™ Mini-PROTEAN® TGX™ precast protein gels (Bio-Rad; Stain-free gels were also used in specific experiments) before electro-transference (Trans-blot Turbo transfer system, Bio-Rad) to nitrocellulose filters. GFP- or HA_3x_-tagged proteins were detected using α-GFP (1:1000; Roche) or α-HA_3x_ (1:1000; Santa Cruz Biotechnology) mouse monoclonal antibody cocktails. Peroxidase-conjugated α-mouse (1:2500, Jackson Immuno-Research Laboratories) was used as secondary antibody. Peroxidase activity was induced using the Clarity Western ECL substrate (Bio-Rad). Chemiluminescence was detected using a Chemidoc + XRS system (Bio-Rad).

### Phosphopeptide enrichment

Aliquots of approximately 300 µg of the crude protein extracts were precipitated with acetone and resuspended in a mixture of urea (8 M) and ammonium bicarbonate (50 mM). Proteins were reduced with dithiotreitol (10 mM) for 30 minutes at room temperature (RT), alkylated with iodoacetamide (50 mM) in the dark for 30 minutes (RT), and digested with 15 µg of trypsin overnight at 37 °C. C18 SEP-PAK columns were used for desalting of peptide mixtures. Samples were dried at RT using a speedvac concentrator.

For TiO_2_ phosphopeptide enrichment, slurry of Titansphere TiO (5 µm) beads (cat. N°.: 5020-75000) was prepared in Buffer C (300 mg mL^-1^ lactic acid; 53 % acetonitrile, ACN; 0.07 % trifluoroacetic acid, TFA) at 25 mg mL^-1^.

Peptides pellets were resuspended in 600 µL of Buffer C, and 72 µL of the Titanium slurry were added to each sample. After incubation for 15-30 minutes at RT with end-over-end rotation, samples were centrifuged for spinning down the beads. Pellets were resuspended in 150 µL of Buffer C and transferred of top of the C8 disks Stage Tips [89]. Buffer C was removed at moderate speed (10-30 µL min^-1^) using a syringe and another 150 µL of Buffer C were added of top of the C8/TiO2 stage tip and passed it through at the same speed for washing. Additional washing was carried out with 100 µL of Buffer B (80 % ACN; 0.1 % TFA) at the same conditions. For sample elution, 2 x 50 µL of Buffer D (0.5 % NH_4_OH) were applied, collecting each eluate in 100 µL of TFA (2 %). Samples were dried in a speedvac concentrator and reconstituted in TFA (0.1 %) for desalting using C 18 Zip-Tip columns.

For mass-spectrometry analysis, fractions of 1/10 from each phosphopeptide-enriched sample were analyzed by nano-LC-MS/MS. Peptides were trapped onto a AcclaimPepMap 100 C18 (2 cm) precolumn (Thermo-Scientific), eluted onto a AcclaimPepMap 100 C18 column (inner diameter 75 μm, 50 cm long, 2 μm particle size; Thermo-Scientific) and separated using a 180 minute gradient (0 - 21 % Buffer B for 60 minutes; 21 % - 35 % Buffer B for 100 minutes, 95 % Buffer B for 10 minutes, and 0 % Buffer B for 10 minutes; Buffer A: 0.1 % formic acid, 2 % ACN; Buffer B: 0.1 % formic acid in ACN) at a flow-rate of 250 nL min^-1^ on a nanoEasy HPLC (Proxeon) coupled to a nanoelectrospray ion source (Thermo-Scientific).

A Q-Exactive mass spectrometer (Thermo-Scientific) in the positive ion mode was used for mass spectra acquisition. Full-scan MS spectra (m/z 400–1,500) were acquired in the Orbitrap at a resolution of 70,000 at m/z 200 and the 10 most intense ions were selected for MS/MS. Fragmentation was carried out with a normalized collision energy of 27 eV. MS/MS scans were acquired with a starting mass of m/z 200; AGC target was 2e^5^, resolution of 17,500 (at m/z 200), intensity threshold of 8e^3^, isolation window of 2.0 m/z units and maximum IT was 100 ms. Charge state screening was enabled to reject unassigned, singly charged, and equal or more than seven protonated ions. A dynamic exclusion time of 20 seconds was used to discriminate against previously selected ions.

Mass spectra *.raw files were searched against the *A. nidulans* fasta database (10,720 protein entries) using Sequest search engine through Proteome Discoverer (v1.4.0.288, Thermo Scientific). Search parameters included a maximum of two missed cleavages allowed, carbamidomethylation of Cys residues as a fixed modification and N-terminal acetylation, C-terminal oxidation and Ser, Thr and Tyr phosphorylation as variable modifications. Precursor and fragment mass tolerance were set to 10 ppm and 0.02 Da, respectively. Node 3 phosphoRS was used as scoring algorithm. This node evaluates statistical confidence of location of phosphorylation sites. Identified peptides were validated using Percolator algorithm with a q-value threshold ≤ 0.01 [90].

### Pull-down assays

Pierce Anti-HA agarose beads (50 µL per sample; ThermoScientific) were centrifuged 5-10 seconds at 12,000 g and washed with a 1:1 volume of TBS (tris buffered saline; ThermoScientific) by centrifugation at the same conditions. After discarding the liquid, 2 mg of crude protein lysate were added. Samples were incubated for 90 minutes at 4 °C with gentle end-over-end mixing. After pelleting the resin with a 5-10 seconds pulse at 12,000 g, supernatants were collected and 200 mg of protein were precipitated with TCA. This fraction was saved for analysis of binding efficiency (non-retained, NR, fraction). Then, the resin was washed thrice (pulses of 5-10 seconds at 12,000 g) with 500 µL of TBS-T (TBS with 0.05 % tween 20). For elution of the bait protein, 1:1 ratios (v/v) of non-reducing 2x SDS-PAGE loading buffer (20 % glycerol, 4 % SDS, 0.004 % bromophenol blue, 12.5 mM tris-HCl, pH = 6,8) were added before incubation for 10 minutes at 30 °C. The resin was pelleted by centrifugation at 12,000 g for 5-10 seconds and the eluate collected (retained fraction, R). Elution was repeated twice. Finally, a 1:1 volume of SDS-PAGE loading buffer was added to the resin, followed by incubation for 5 minutes at 95 °C. Both NR and R fractions were resolved by SDS-PAGE electrophoresis and analyzed by immunodetection. R fractions were then analyzed by LC-MS/MS.

GFP-Trap beads (25 µL per sample; Proteintech) were diluted in 500 µL of dilution buffer (5 mM HEPES pH = 7.5, 0.5 mM EDTA, 0.1 % NP-40, 20 mM KCl, 150 mM NaCl, plus a protease inhibitor cocktail from Roche) and centrifuged for 2 minutes and 2,500 g at 4 °C. This washing step was repeated twice. Then, 3 mg of the crude protein extract containing the bait chimera (FlpA::GFP) were added and samples were incubated with gentle end-over-end mixing at 4 °C for 90 minutes. Samples were centrifuged for 2 minutes and 2,500 g at 4 °C. The supernatants were collected and 200 µg of this fraction were precipitated with TCA (fraction NR1). The resin was washed twice with 500 µL of dilution buffer. Then, 3 mg of crude protein extracts containing the Stk47::HA_3x_ chimera were added, and the procedure was repeated (fraction NR2). Finally, 75 µL of SDS-PAGE loading buffer were added, samples were incubated at 95 °C for 5 minutes and centrifuged for 2 minutes and 2,500 g at 4 °C (fraction R). Both NR and R fractions were resolved by SDS-PAGE electrophoresis and analyzed by immunodetection.

## Author contribution

Conceptualization, O.E, X.G. and E.A.E; investigation, Z.A., A.O., A.F., X.G, A.M.-V., E.A.E. and O.E; methodology, E.A.E., X.G., Z.A. and O.E.; supervision, E.A.E, X.G., O.E.; writing—original draft, Z.A. and O.E.; review and editing, Z.A., A.O., A.F., X.G., A.M.-V., J.R.F., E.A.E. and O.E.

## Acknowledgements.

We want to thank Dr. Ricardo Andrade, from the central microscopy services of the University of the Basque Country, for the assistance with their fluorescence microscope. The Δ*stk47::riboB* strain of *A. nidulans* was generated by E. Requena, currently a PhD student at the Instituto Nacional de Investigación y Tecnología Agroalimentaria (INIA; Madrid, Spain).

## Supplementary material

**S1 Table. Probable FlpA, Stk47 and FlpB orthologs identified by BLAST analyses.** The table shows the accession number of each sequence, the taxonomy of the species each sequence belongs to, the length as well as score, expect and coverage values. It also shows the results (first hit) of the confirmatory reverse retrieval of each sequence against the *Aspergillus nidulans* proteome in the AspGD database. The color key used is shown on the right of the table and applies to classes of fungi.

**S2 Table. List of phosphopeptides and phosphorylated proteins identified in crude protein extracts of the wild-type and the null *flpA* or null *stk47* strains.**

**S3 Table. Strains used in this work**

**S4 Table. Oligonucleotides used in this study.**

**S1 File. Candidate exonic mutations found in FLIP57 and FLIP76.** Exonic mutations in FLIP57 (track 1) and FLIP76 (track 2) genomes are shown in comparison to the FGSCA4 reference genome, FLIP166 genome (track 3) or the transcriptomes of wild-type (track 4) or Δ*flbB* (track 5) strains [24,30,91].

**S1 Figure. Phenotypes of selected FLIP mutants.** Images were taken after 72 hours of culture at 37 °C on AMM (row 1) or AMM supplemented with 0.65 M NaH_2_PO_4_ (row 2). Parental Δ*flbB* and FLIP166 mutants were used as controls. Diameter of plates: 5.5 cm.

**S2 Figure. Predicted three-dimensional structures of FlpA (A), Stk47 (B) and FlpB (C), modelled by Swiss-Model.** The models cover the regions His26-Lys374 of FlpA, Ser746-Leu1052 in the case of Stk47 and Glu7-Asn234 in the case of FlpB, and are based on PDB structure 7jv7.1 [36], which can be seen in panel D. The alignments between query sequences and the reference structure can be seen below Swiss-Models.

**S3 Figure: Subcellular localization of FlpA::GFP** when it is expressed driven by its native promoter, in vegetative of A) a wild-type strain, or B) an HhoA::mCherry strain. Scale bars = 5 µm.

**S4 Figure: Phosphopeptide-enrichment analysis for A. nidulans wild-type,** Δ***flpA* and** Δ***stk47* strain**. A) Venn diagram showing the number of phosphorylated proteins detected in wild-type, Δ*flpA* and Δ*stk47* strains. The subgroups of phosphoproteins not detected in the null *flpA* background are highlighted. B) Bar-graph showing the most represented InterPro functions in the list of phosphoproteins not detected in the null *flpA* background compared to the parental wild-type strain. AN8190/Stk47 is included among the ten serine/threonine-protein kinases lost in the Δ*flpA* strain.

**S5 Figure: Enrichment of Stk47::HA_3x_ or FlpA::GFP using HA-agarose resin (Pierce).** Crude protein extracts of strains expressing Stk47::HA_3x_ (wild-type, Δ*flpA* or Δ*flpB* genetic backgrounds) or FlpA::GFP (wild-type background) chimeras were incubated with the resin. Retained fractions were sequentially eluted using non-reducing Laemli buffer (R1, R2 and R3) or SDS-PAGE loading buffer (Rdes). See Materials and Methods. LC-MS/MS analyses of those R fractions did not show significant enrichment in peptides corresponding to putative interactors (FlpA, Stk47 and FlpB) or any other *A. nidulans* cyclin and/or kinases (not shown).

**S6 Figure: Phenotype of *A. fumigatus* single-null mutants of *flpA*, *flpB* and *stk47*** on AMM medium, under oxygen- or iron-limiting conditions, or under oxidative stress, after 48 hours of growth at 37 °C. Growth inhibition halos were measured after 48 hours.

**S7 Figure: Susceptibility of single-null *flpA*, *flpB* and *stk47* strains of *A. fumigatus* to posaconazole, voriconazole and caspofungin**, determined using gradient diffusion strips in RPMI agar plates. Pictures correspond to 48 hours of growth at 37 °C.

