## Supplementary figures and images for "Identification and functional characterization of the putative members of the CTDK-1 kinase complex as regulators of growth and development in the genus *Aspergillus*"

### Figure S1

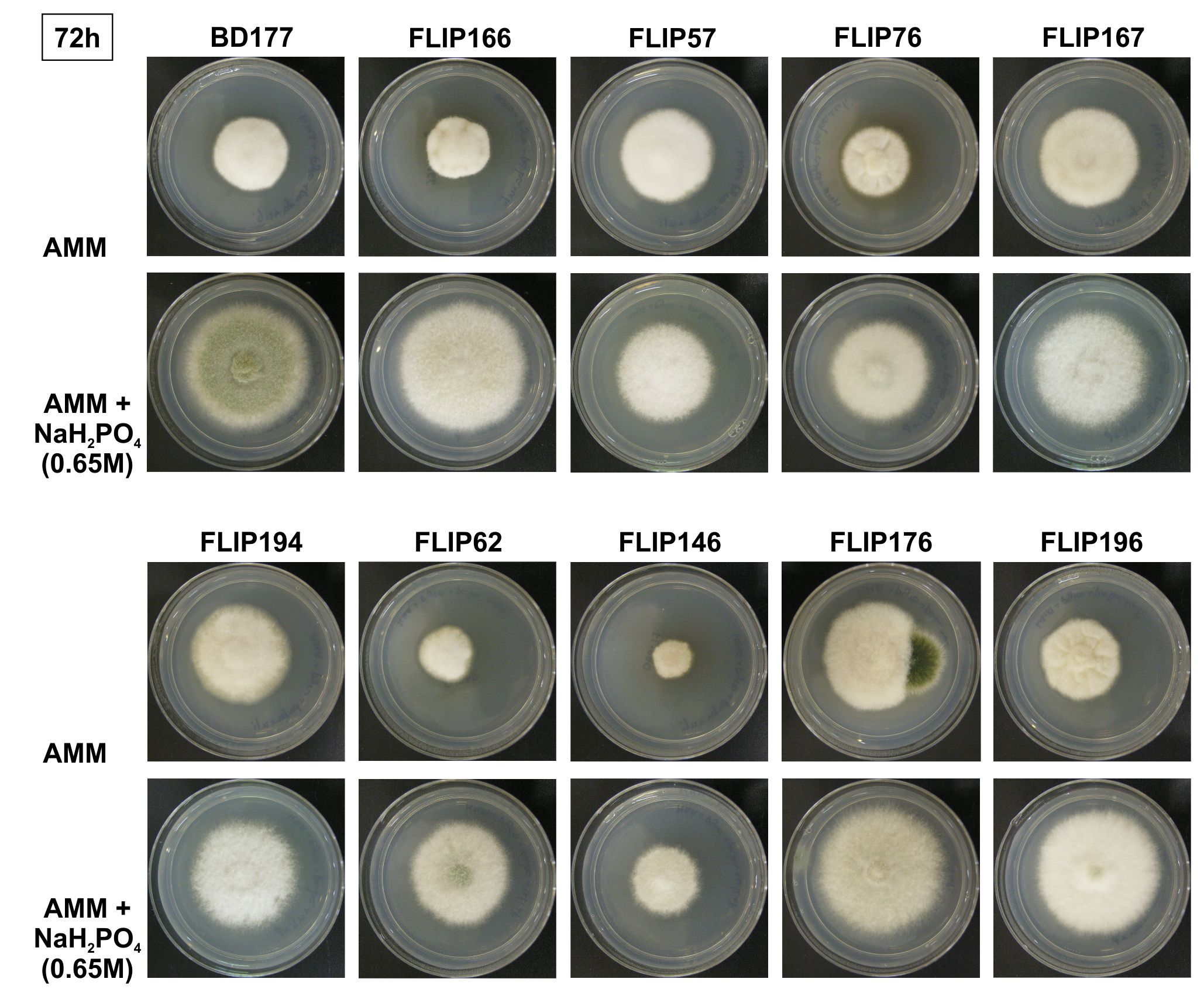

### Figure S2

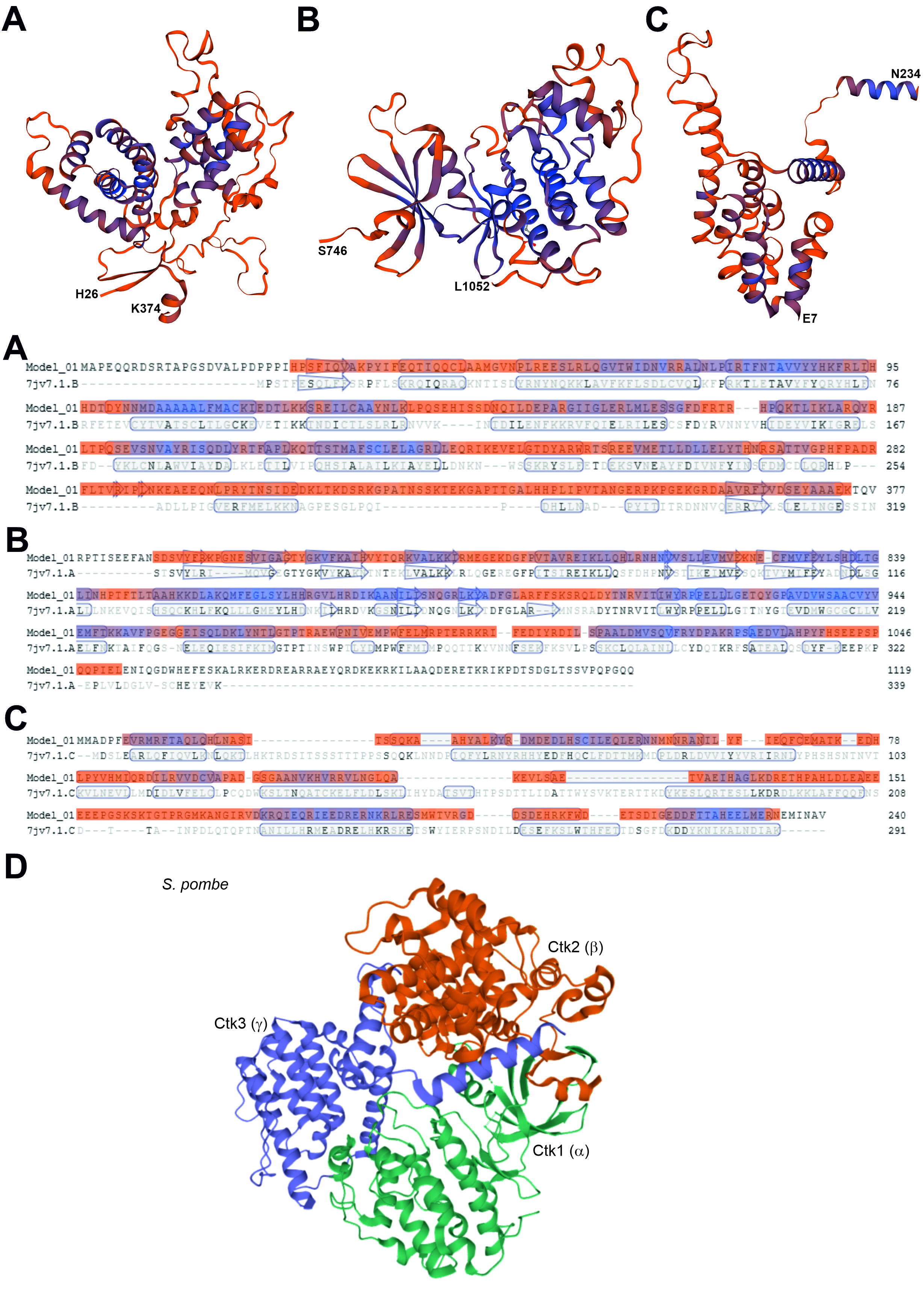

### Figure S3

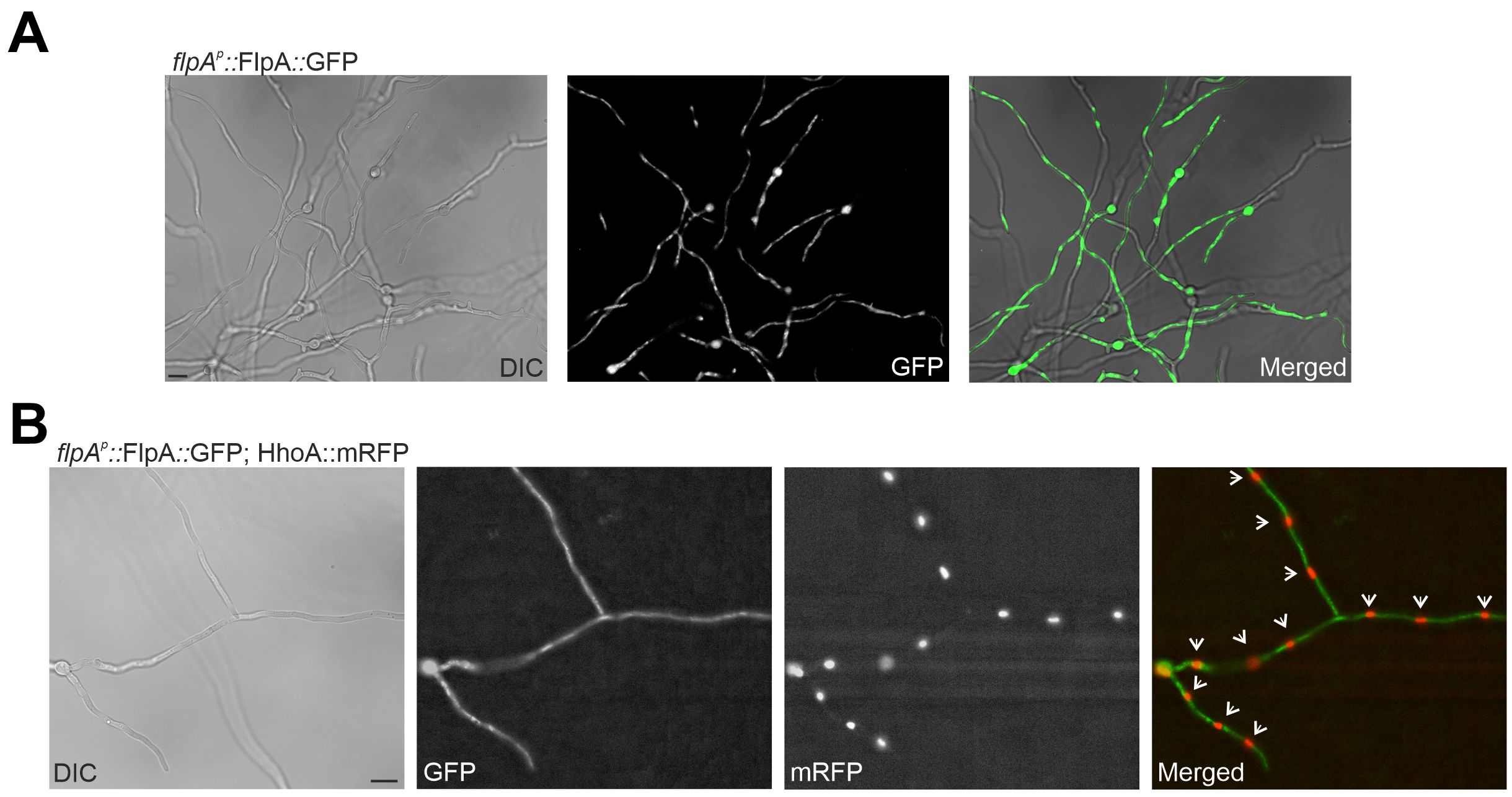

### Figure S4

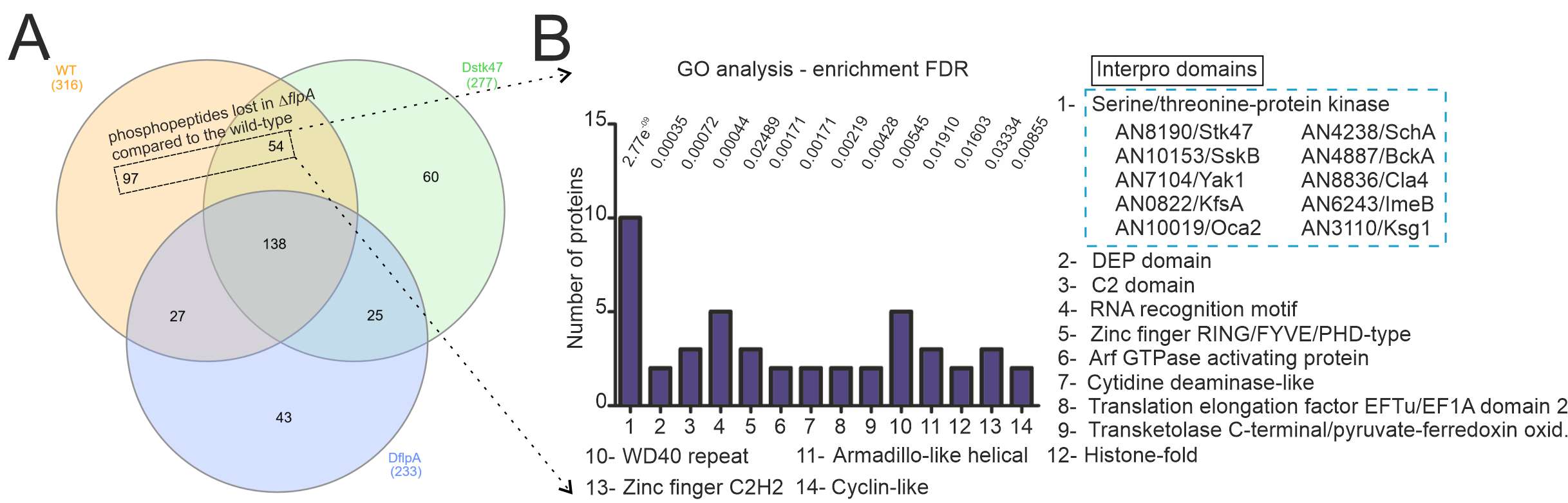

### Figure S5

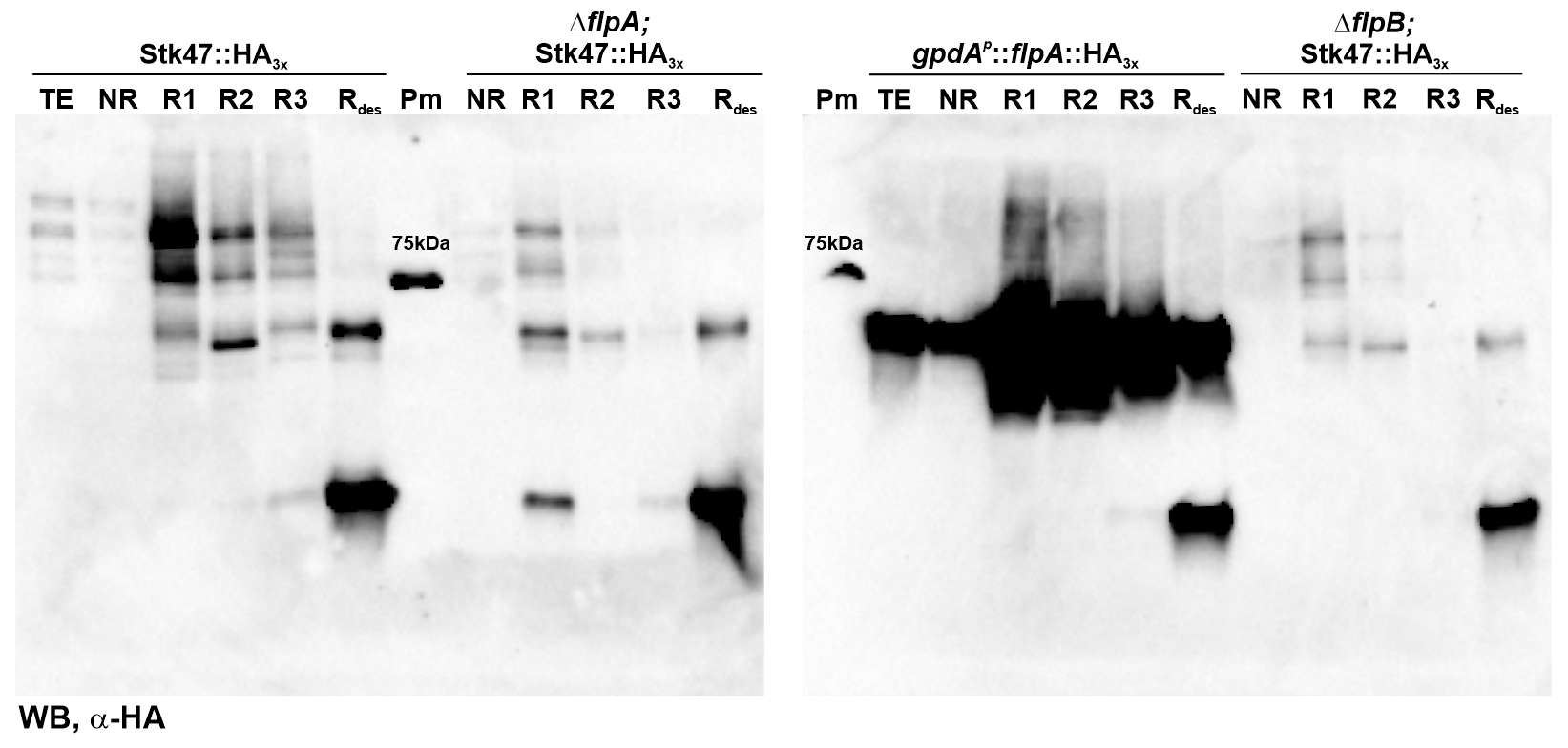

### Figure S6

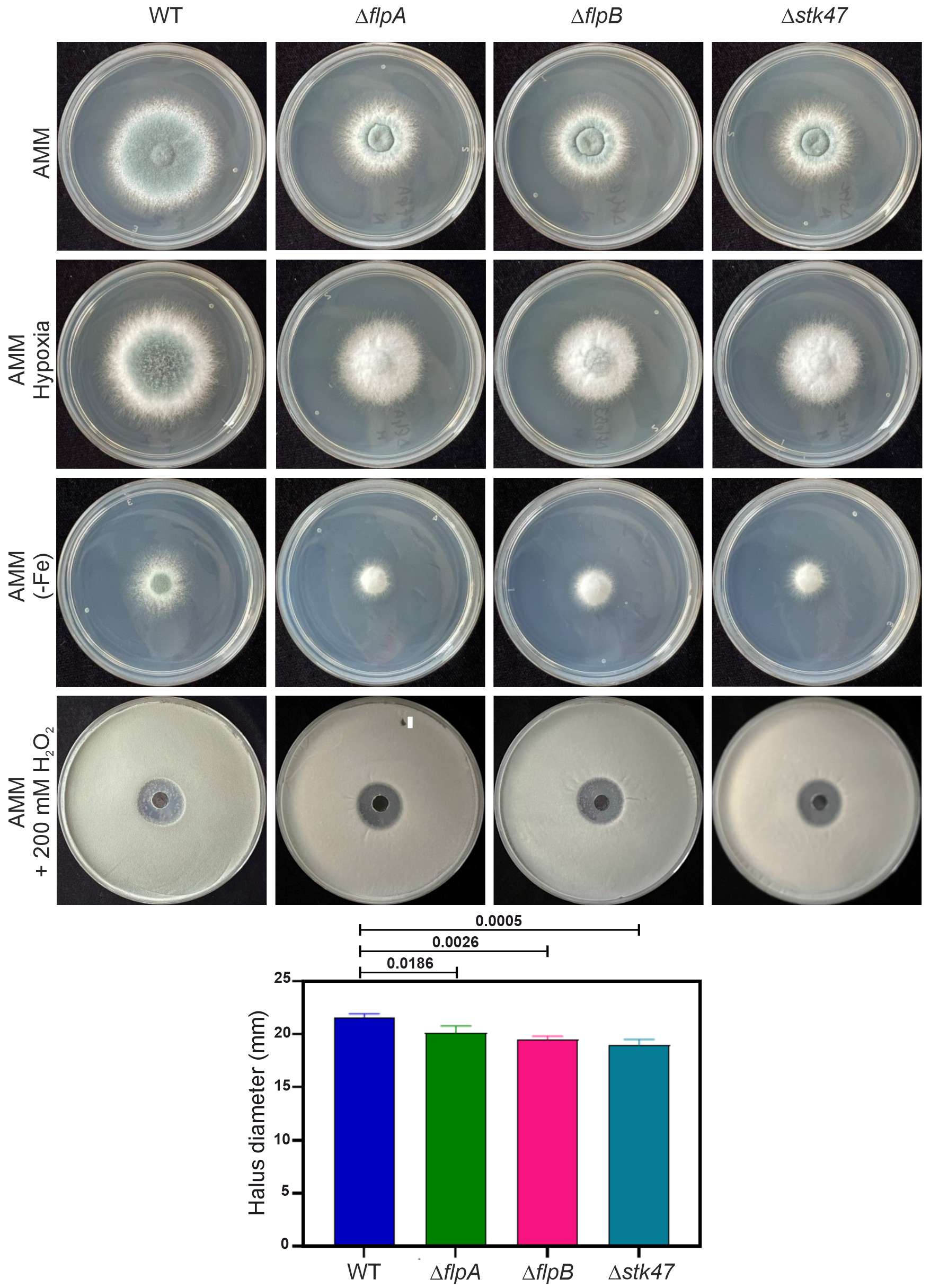

### Figure S7

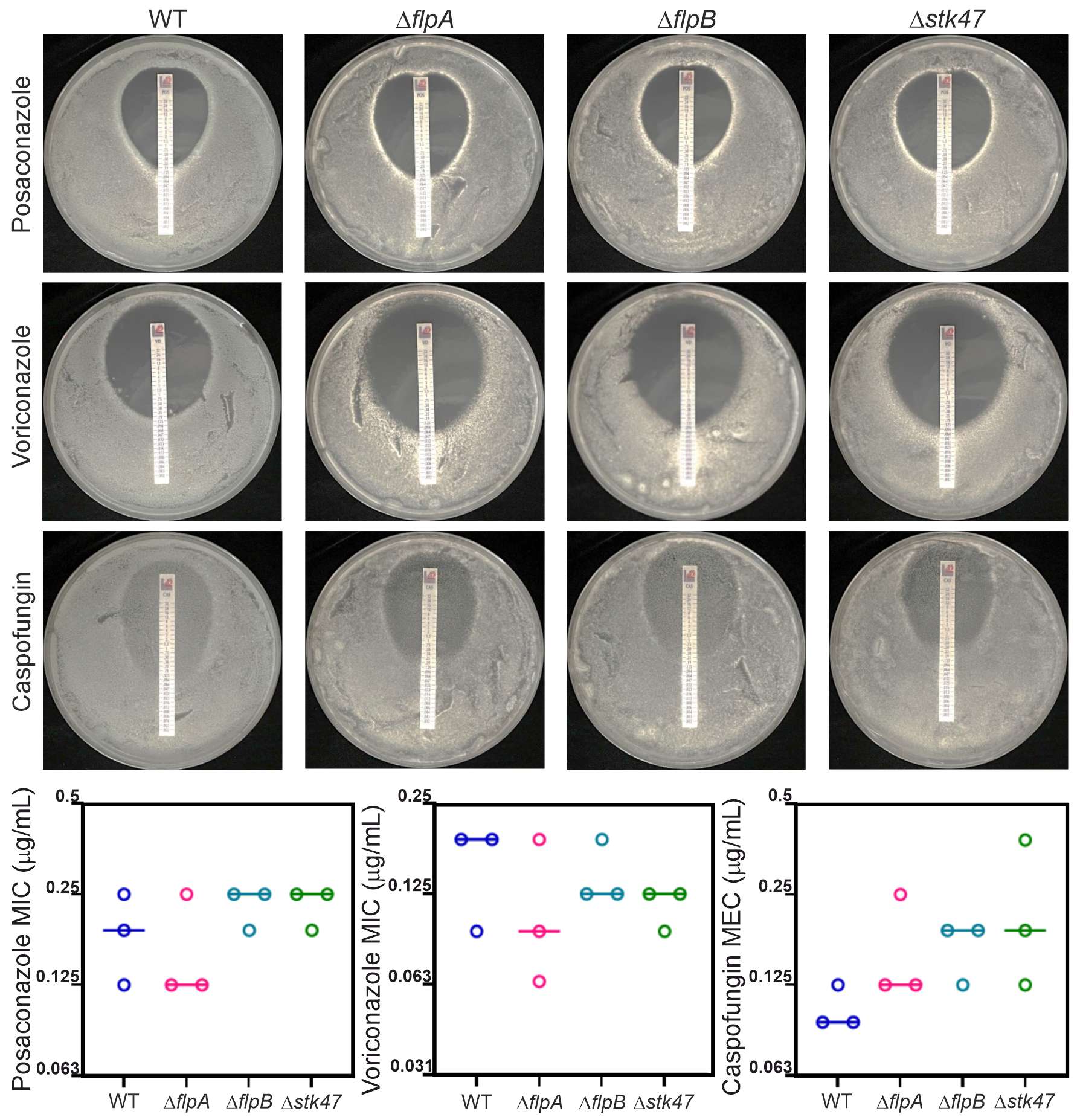
