## Supplementary material for "Identification and functional characterization of the putative members of the CTDK-1 kinase complex as regulators of growth and development in the genus *Aspergillus*": File S1

FLIP57:

1) An6932 (ChrI; 3620023 bp): High-capacity, high-affinity uric acid-xanthine permease; induced by purine; positively regulated by uaY; under ammonium repression by AreA; localized to plasma membrane and strongly expressed in periphery of metulae.

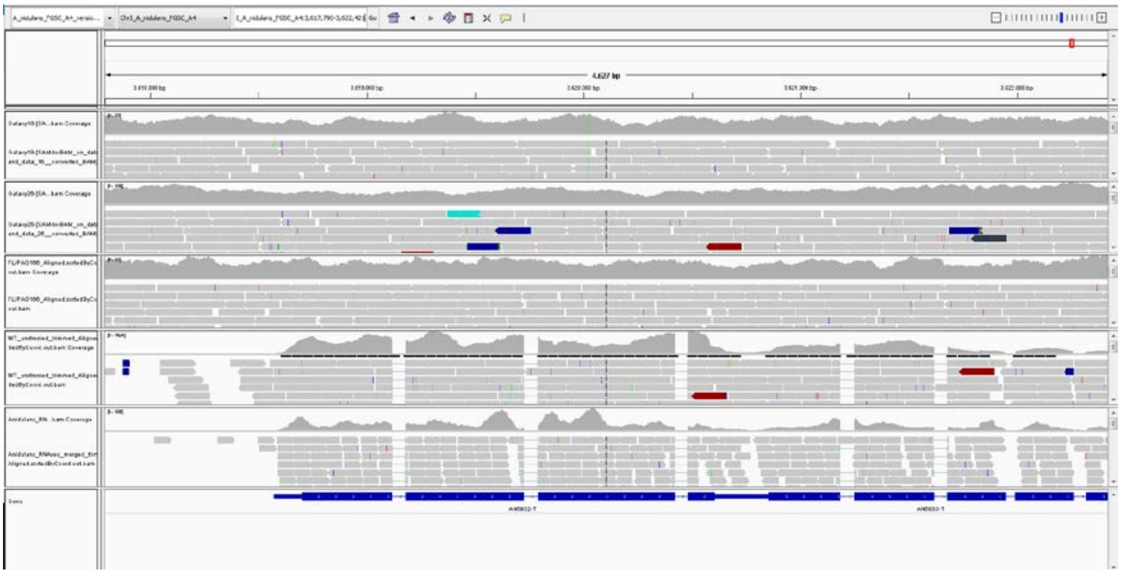

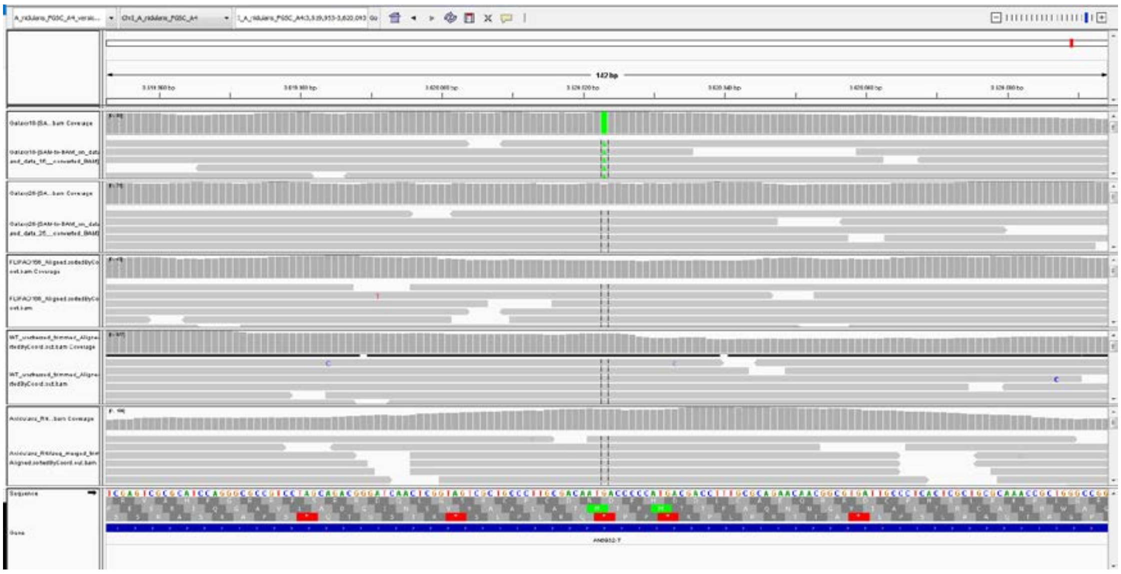

Met (ATG) in position 400 by an Ile (ATA).

2) An7856 (ChrII; 22962 bp): Has domain(s) with predicted enoyl-[acyl-carrier-protein] reductase (NADH) activity, fatty acid synthase activity, role in fatty acid biosynthetic process, oxidation-reduction process and fatty acid synthase complex localization

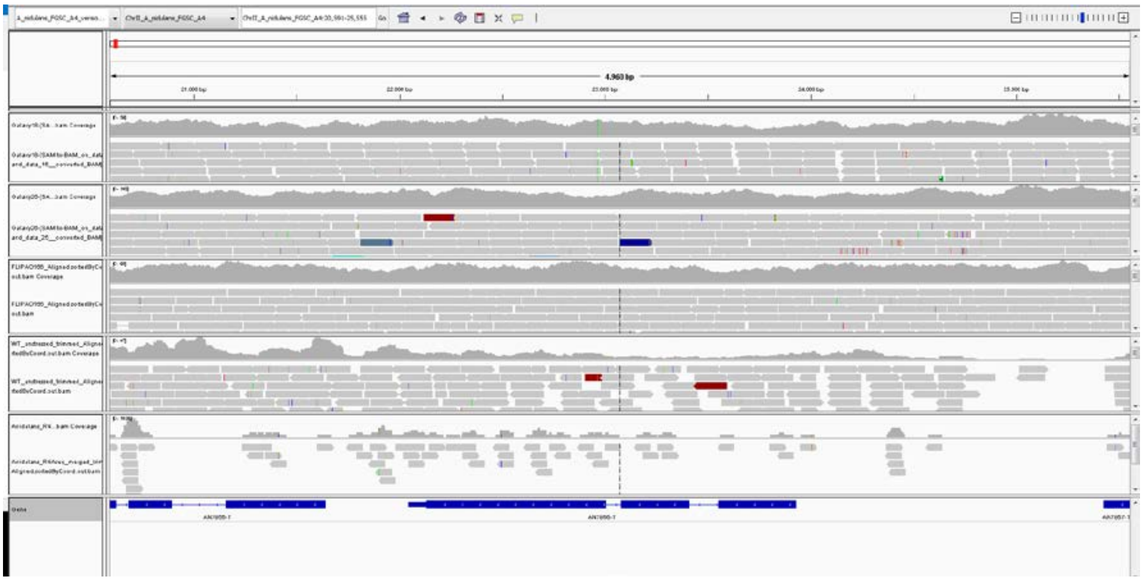

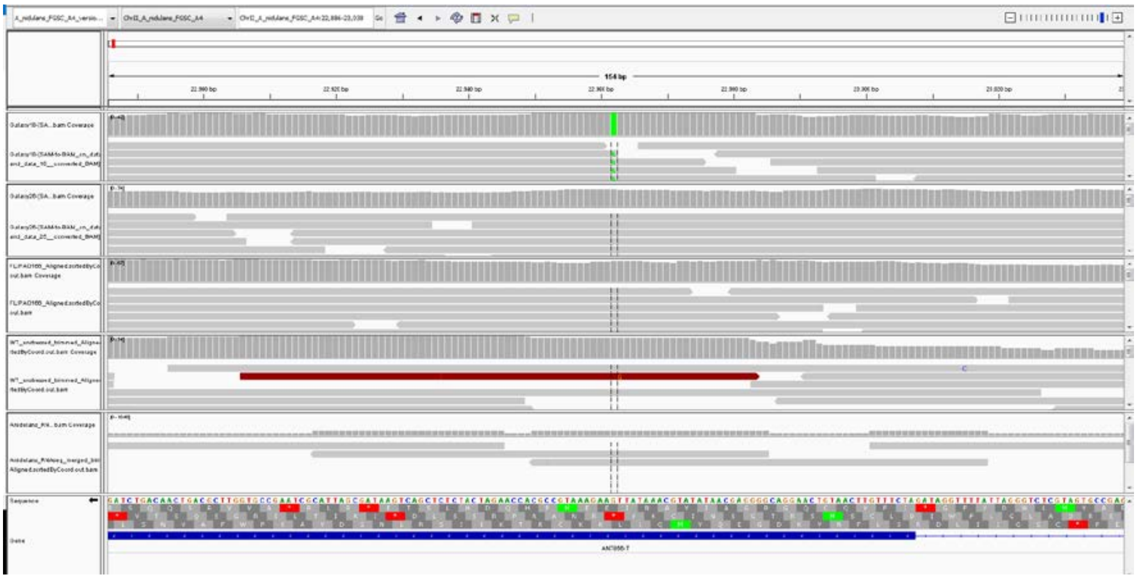

Leu (TTG) in position 253 by a Phe (TTT).

3) An7842 (ChrIV; 2854010 bp): Protein of unknown function.

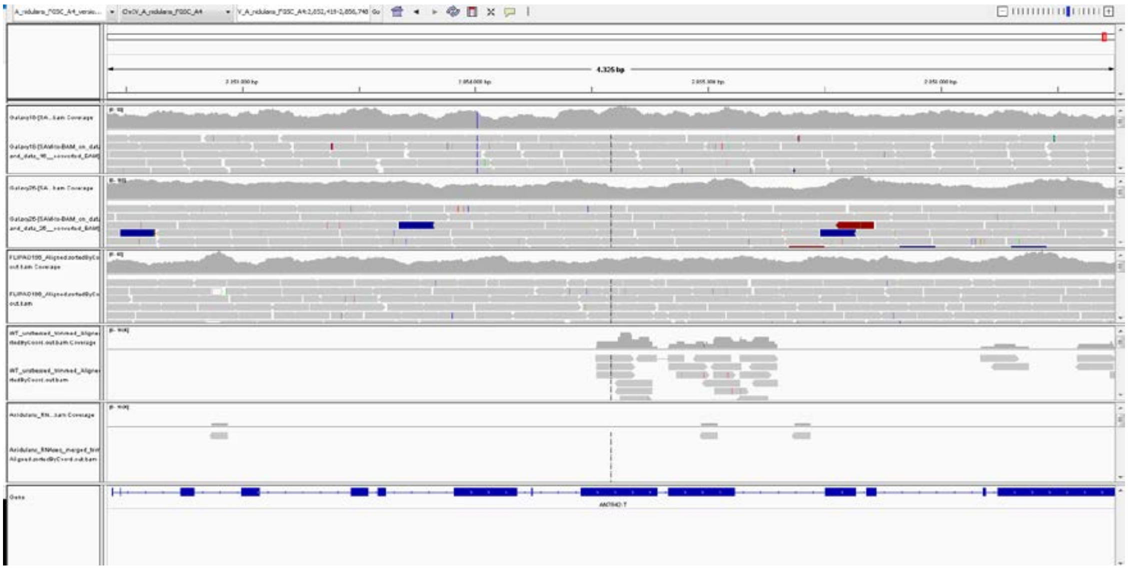

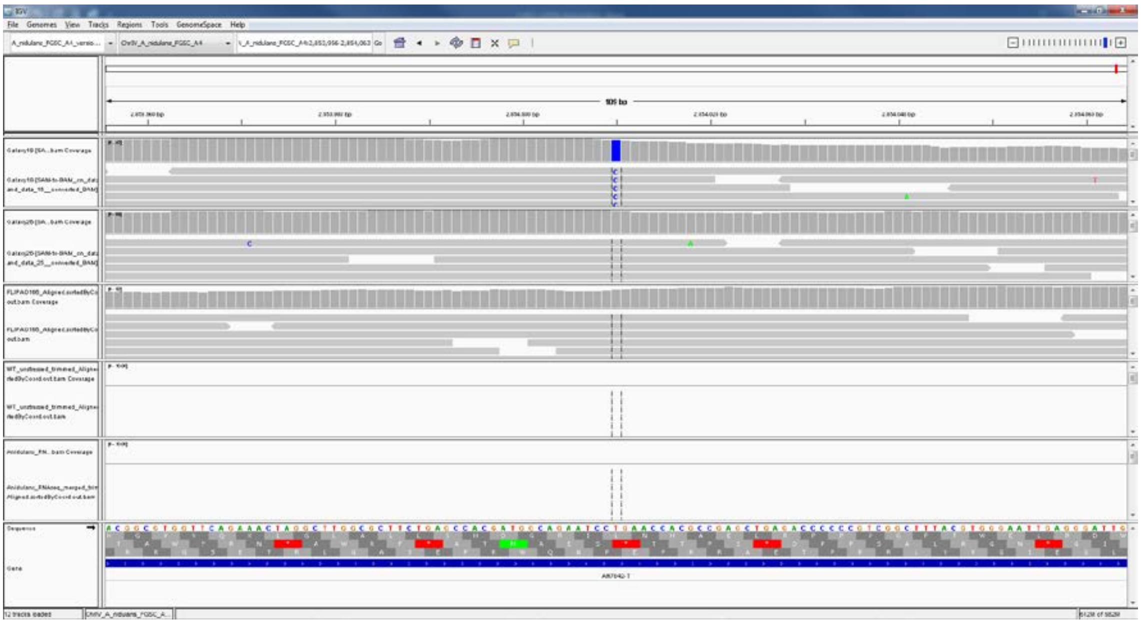

Leu (CTG) in position 120 by a Pro (CCG).

4) An10640 (Chr V; 1324727 bp): Ortholog(s) have cyclin-dependent protein serine/threonine kinase activator activity

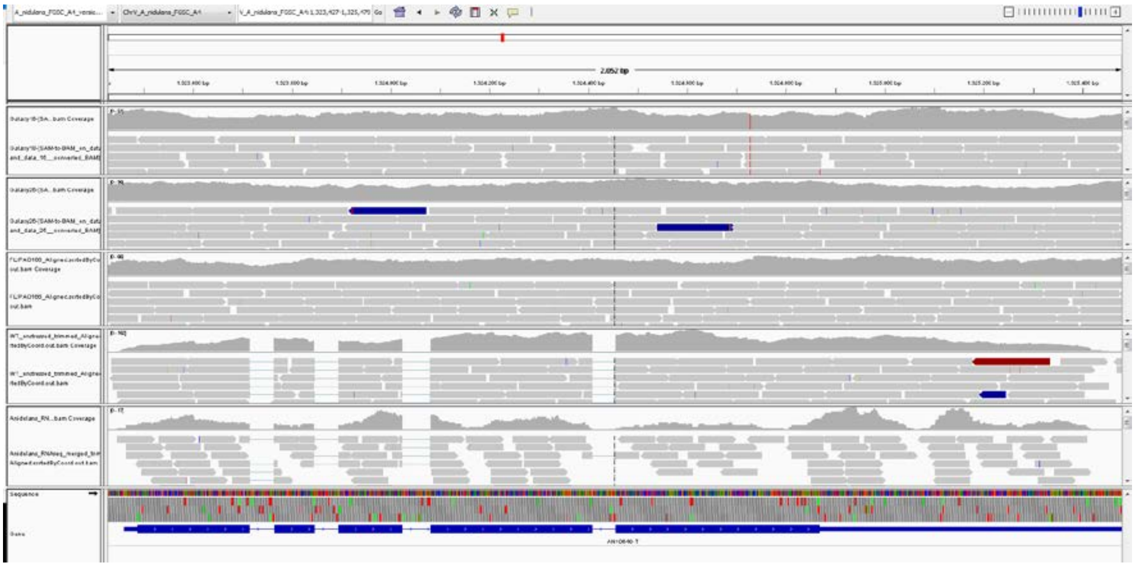

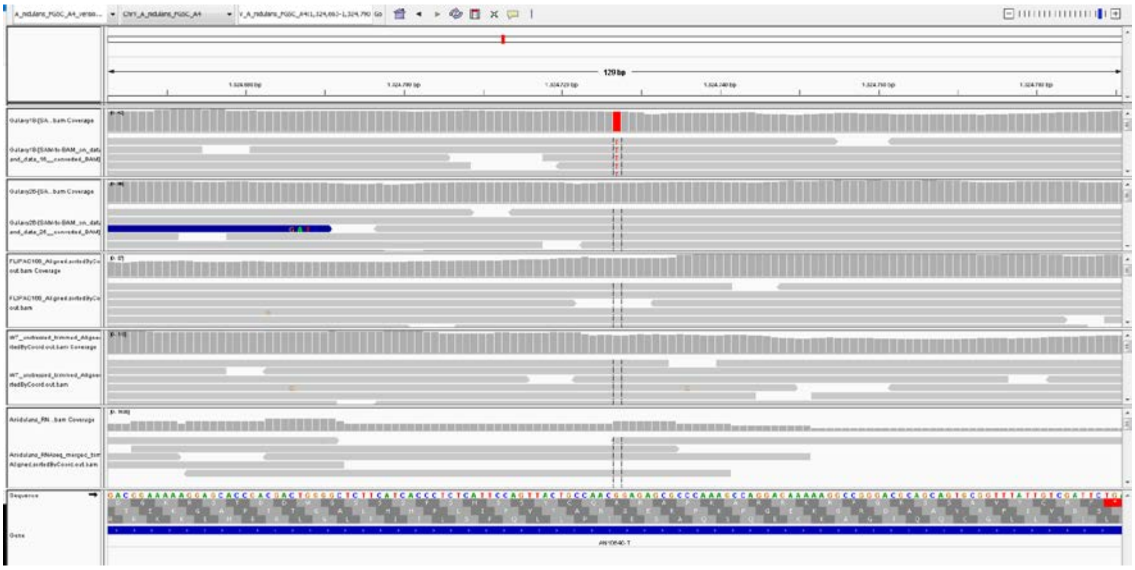

Gly (GGA) in position 347 by a stop codon (TGA)

FLIP76:

1) An12237/abpA (ChrVII; 598883 bp): Putative actin-binding protein of the cortical actin patches involved in endocytosis

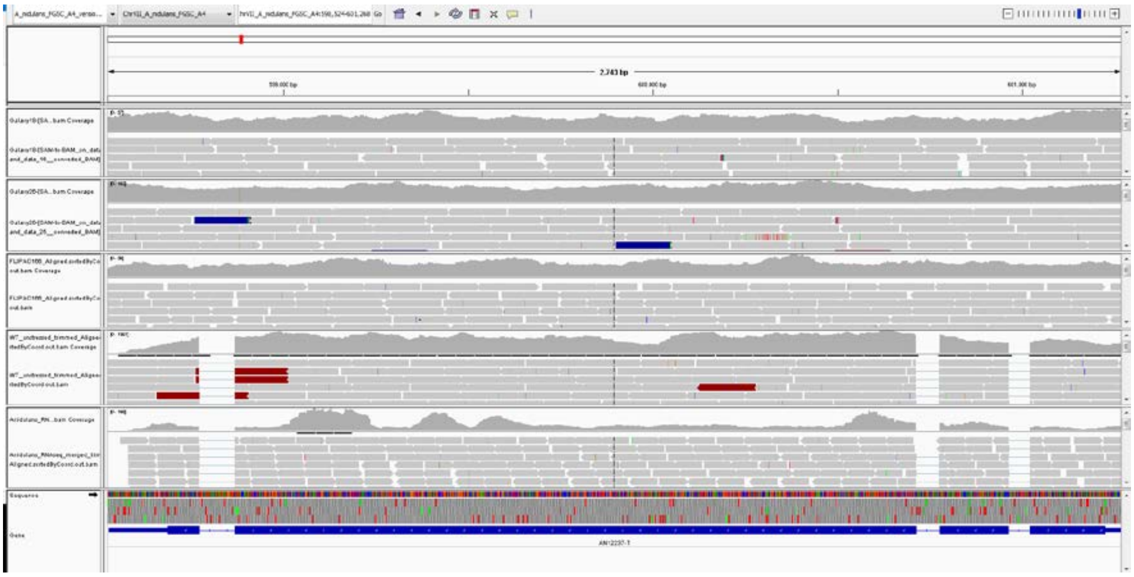

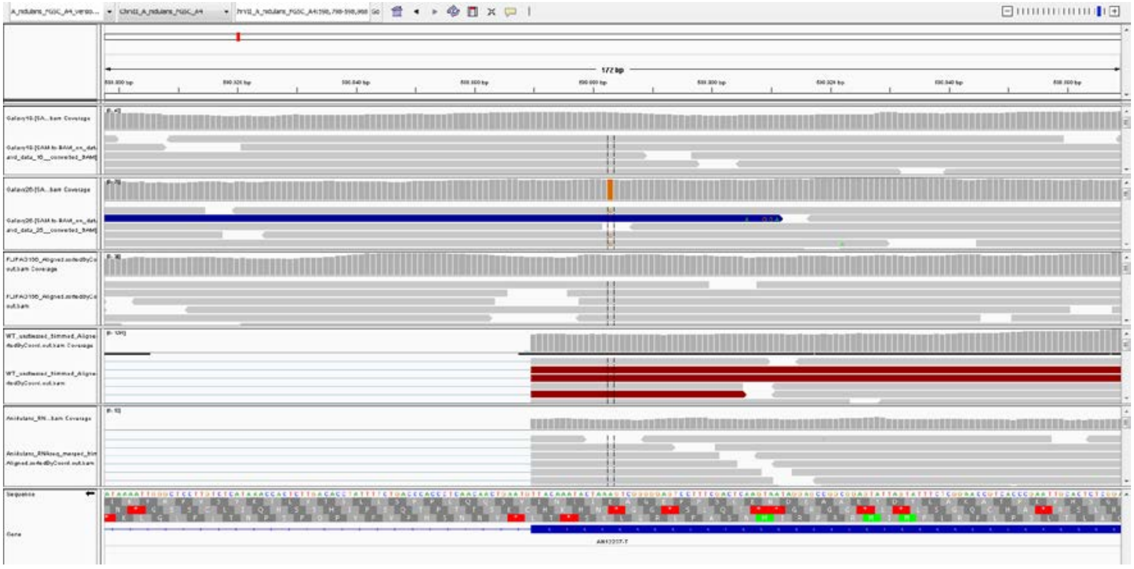

Glu (GAA) in position 742 by an Ala (GCA).

2) An5221 (ChrV; 1537739 bp): ORF uncharacterized.

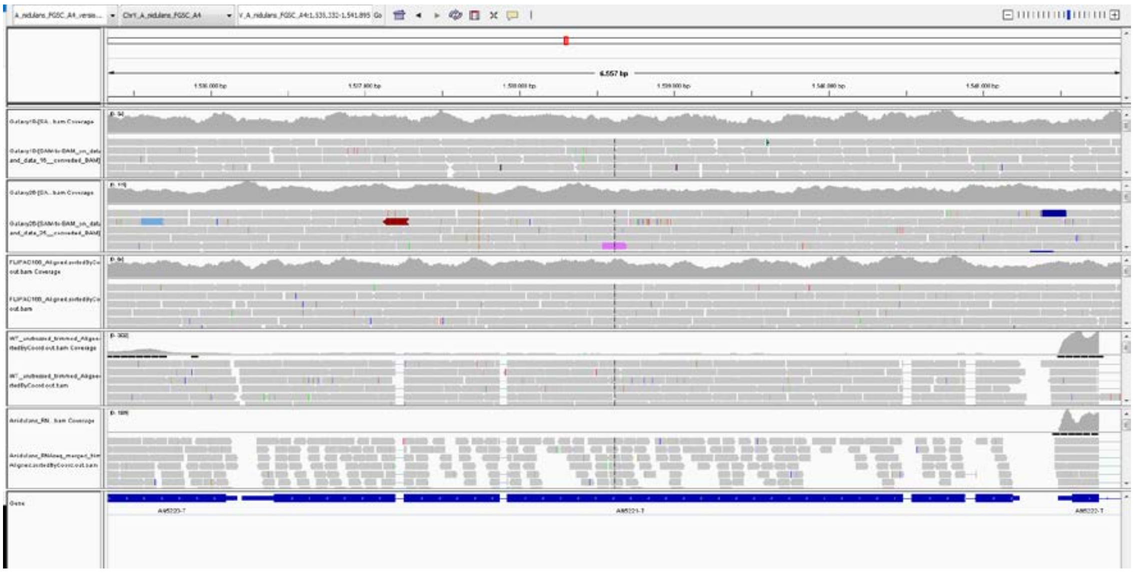

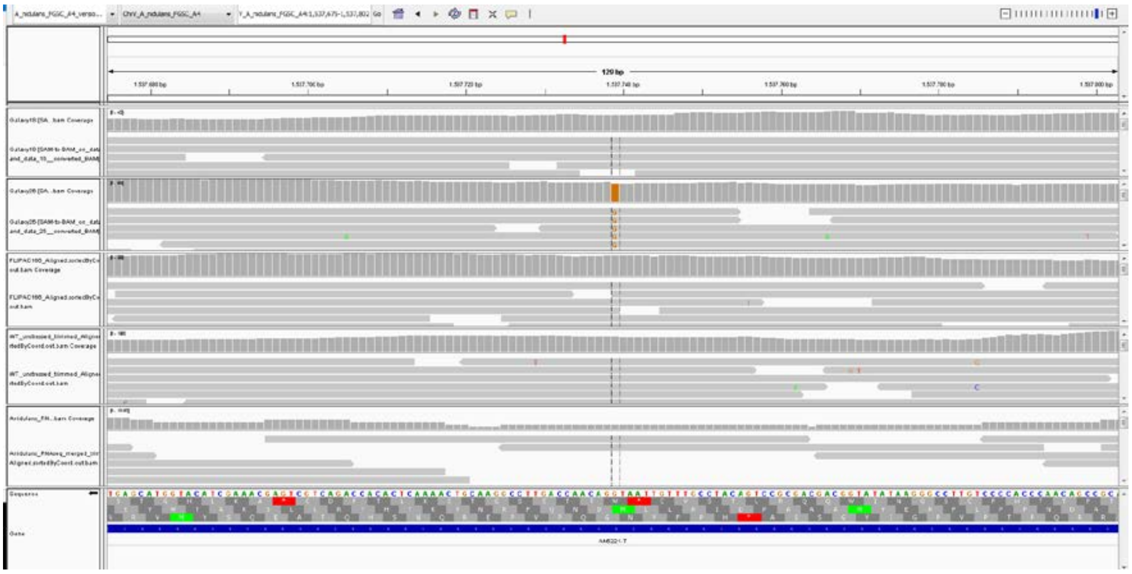

Met (ATG) in position 1097 by an Ile (ATC).

3) An5717 (ChrV; 1901598 bp): Non-essential karyopherin family protein; required for normal hyphal growth and conidial development.

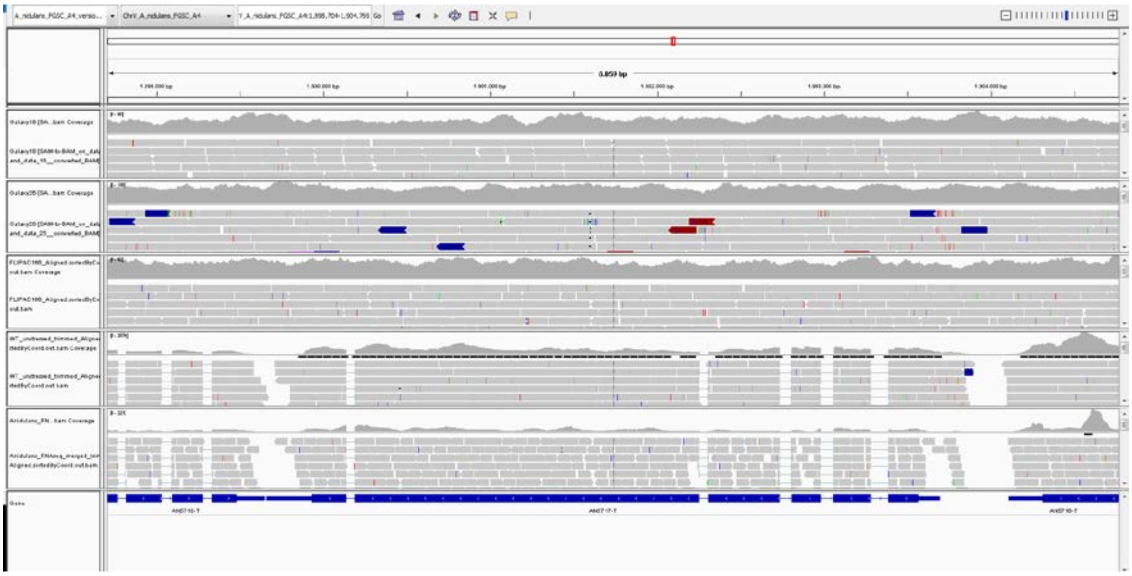

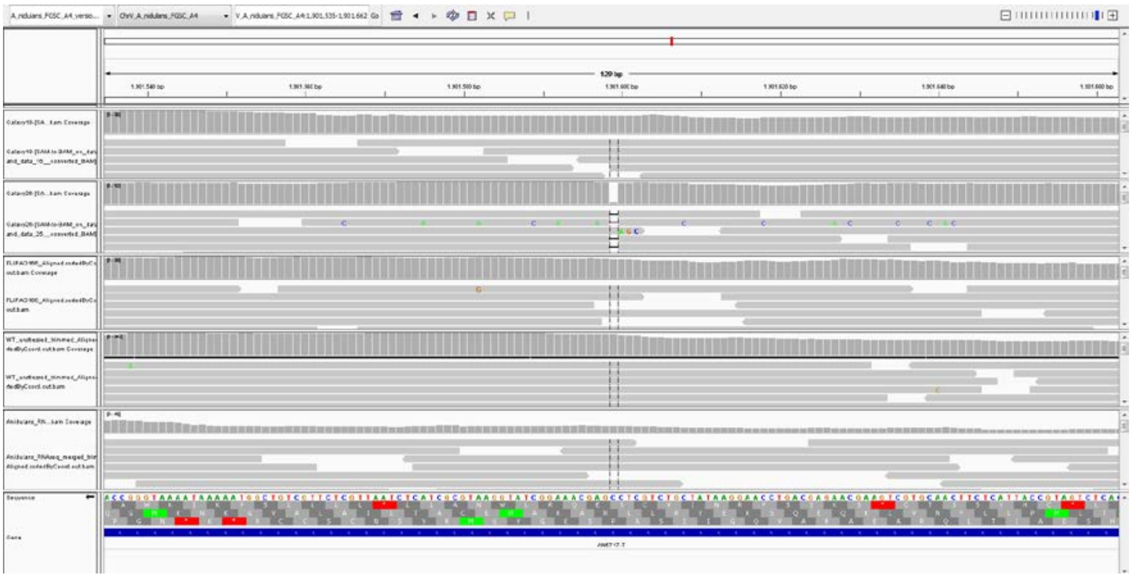

Arg (CGA) in position 557 by EQRLWNALL-Stop (Arg557Glu+8-Stop).
