## Supplementary material for "Identification and functional characterization of the putative members of the CTDK-1 kinase complex as regulators of growth and development in the genus *Aspergillus*": Table S3

**Table S3. Strains used in this work**

| **Name** | **Genotype** | **Reference** |
| --- | --- | --- |
| ***Aspergillus nidulans* strains** | | |
| BD177 | *pyrG89; argB2;* Δ*flbB::riboB^Afum^, pabaB22, pyroA4,* Δ*nkuA::argB; riboB2, veA1* | (Garzia *et al.*, 2009) |
| BD950 (FLIP166) | *pyrG89; An12172^Ile94Asn^; argB2;* Δ*flbB::riboB^Afum^, pabaB22, pyroA4,* Δ*nkuA::argB; pmtC^Pro282Leu^, acdA^Leu99Arg^;riboB2, veA1* | (Otamendi *et al.*, 2019a) |
| BD1339 (FLIP57) | *pyrG89; argB2;* Δ*flbB::riboB^Afum^, pabaB22, pyroA4,* Δ*nkuA::argB; flpA^Gly347Stop^; riboB2, veA1* | (Otamendi *et al.*, 2019a) |
| BD1341 (FLIP76) | *pyrG89; argB2;* Δ*flbB::riboB^Afum^, pabaB22, pyroA4,* Δ*nkuA::argB; kapI^Arg557Glu^; riboB2, veA1* | (Otamendi *et al.*, 2019a) |
| BD1373 | *pyrG89; argB2; pyroA4,* Δ*nkuA::argB;* Δ*flpA::pyrG^Afum^*; *veA1* | This study |
| BD1374 | *pyrG89; argB2; pyroA4,* Δ*nkuA::argB; flpA::ha_3x_::pyrG^Afum^*; *veA1* | This study |
| BD1376 | *pyrG89; argB2; pyroA4,* Δ*nkuA::argB; flpA^Gly347Stop^::ha_3x_::pyrG^Afum^*; *veA1* | This study |
| BD1378 | *pyrG89; argB2; pyroA4,* Δ*nkuA::argB; flpA::gfp::pyrG^Afum^*; *veA1* | This study |
| BD1384 | *pyrG89; argB2;* Δ*flbB::riboB^Afum^, pabaB22, pyroA4,* Δ*nkuA::argB;* Δ*flpA::pyrG^Afum^*; *riboB2, veA1* | This study |
| BD1385 | *pyrG89; argB2;* Δ*flbB::riboB^Afum^, pabaB22, pyroA4,* Δ*nkuA::argB; flpA::ha_3x_::pyrG^Afum^*; *riboB2, veA1* | This study |
| BD1387 | *pyrG89; argB2;* Δ*flbB::riboB^Afum^, pabaB22, pyroA4,* Δ*nkuA::argB; flpA^(Gly347Stop)^::ha_3x_::pyrG^Afum^*; *riboB2, veA1* | This study |
| BD1388 | *pyrG89; argB2;* Δ*flbB::riboB^Afum^, pabaB22, pyroA4,* Δ*nkuA::argB; flpA::gfp::pyrG^Afum^*; *riboB2, veA1* | This study |
| BD1391 | *pyrG89; argB2;* Δ*flbB::riboB^Afum^, pabaB22, pyroA4,* Δ*nkuA::argB; flpA^(Gly347Stop)^::gfp::pyrG^Afum^*; *riboB2, veA1* | This study |
| BD1392  (FLIP57 background) | *pyrG89; argB2;* Δ*flbB::riboB^Afum^, pabaB22, pyroA4,* Δ*nkuA::argB; flpA::ha_3x_::pyrG^Afum^*; *riboB2, veA1* | This study |
| BD1394  (FLIP57 background) | *pyrG89; argB2;* Δ*flbB::riboB^Afum^, pabaB22, pyroA4,* Δ*nkuA::argB; flpA::gfp::pyrG^Afum^*; *riboB2, veA1* | This study |
| BD1397  (FLIP57 background) | *pyrG89; argB2;* Δ*flbB::riboB^Afum^, pabaB22, pyroA4,* Δ*nkuA::argB; flpA^(Gly347Stop)^::gfp::pyrG^Afum^*; *riboB2, veA1* | This study |
| BD1438 | *pyrG89; argB2; pyroA4,* Δ*nkuA::argB; flpA::gfp::pyrG^Afum^*; *hhoA::mCh::pyroA^Afum^; veA1* | This study |
| BD1442 | *pyrG89; argB2; pyroA4,* Δ*nkuA::argB; flpA^p^::gpdA^p^::flpA::gfp::pyrG^Afum^*; *veA1* | This study |
| BD1446 | *pyrG89;* Δ*stk47::pyrG^Afum^*; *argB2; pyroA4,* Δ*nkuA::argB; veA1* | This study |
| BD1448 | *pyrG89; stk47::ha_3x_::pyrG^Afum^*; *argB2; pyroA4,* Δ*nkuA::argB; veA1* | This study |
| BD1451 | *pyrG89; stk47::gfp::pyrG^Afum^*; *argB2; pyroA4,* Δ*nkuA::argB; veA1* | This study |
| BD1454 | *pyrG89,* Δ*flpB::pyrG^Afum^*; *argB2; pyroA4,* Δ*nkuA::argB; veA1* | This study |
| BD1457 | *pyrG89, flpB::gfp::pyrG^Afum^*; *argB2; pyroA4,* Δ*nkuA::argB; veA1* | This study |
| BD1466 | *pyrG89; stk47::gfp::pyroA^Afum^*; *argB2; pyroA4,* Δ*nkuA::argB;* Δ*flpA::pyrG^Afum^*; *veA1* | This study |
| BD1468 | *pyrG89, flpB::gfp::pyroA^Afum^; argB2; pyroA4,* Δ*nkuA::argB;* Δ*flpA::pyrG^Afum^*; *veA1* | This study |
| BD1473 | *pyrG89,* Δ*flpB::pyroA^Afum^;argB2; pyroA4,* Δ*nkuA::argB;* Δ*flpA::pyrG^Afum^*; *veA1* | This study |
| BD1479 | *pyrG89,* Δ*flpB::pyrG^Afum^*; *stk47::gfp::pyroA^Afum^*; *argB2; pyroA4,* Δ*nkuA::argB; veA1* | This study |
| BD1485 | *pyrG89,* Δ*flpB::pyrG^Afum^*; *argB2; pyroA4,* Δ*nkuA::argB; flpA^p^::gpdA^p^::flpA::gfp::pyroA^Afum^*; *veA1* | This study |
| BD1487 | *pyrG89;* Δ*stk47::riboB^Afum^*; *argB2; pabaB22;* Δ*nkuA::argB;* Δ*flpA::pyrG^Afum^*; *riboB2, veA1* | This study |
| BD1494 | *pyrG89;* Δ*stk47::riboB^Afum^*; *argB2; pabaB22;* Δ*nkuA::argB; flpA^p^::gpdA^p^::flpA::gfp::pyrG^Afum^*; *riboB2, veA1* | This study |
| BD1495 | *pyrG89,* Δ*flpB::pyrG^Afum^;* Δ*stk47::riboB^Afum^*; *argB2; pabaB22;* Δ*nkuA::argB; riboB2, veA1* | This study |
| BD1499 | *pyrG89, flpB::gfp::pyrG^Afum^;* Δ*stk47::riboB^Afum^*; *argB2; pabaB22;* Δ*nkuA::argB; riboB2, veA1* | This study |
| BD1503 | *pyrG89; stk47::ha_3x_::pyrG^Afum^*; *argB2; pyroA4,* Δ*nkuA::argB;* Δ*flpA::pyroA^Afum^*; *veA1* | This study |
| BD1506 | *pyrG89,* Δ*flpB::pyroA^Afum^; stk47::ha_3x_::pyrG^Afum^*; *argB2; pyroA4,* Δ*nkuA::argB;*; *veA1* | This study |
| MAD6500 | *pyrG89;* Δ*stk47::riboB^Afum^*; *argB2; pabaB22;* Δ*nkuA::argB; riboB2, veA1* | Elena Requena-Eduardo A. Espeso |
| TN02A3 | *pyrG89; argB2; pyroA4,* Δ*nkuA::argB; veA1* | (Nayak *et al.*, 2006) |
| ***Aspergillus fumigatus* strains** | | |
| ΔakuB::pyrG |  | (Al Abdallah *et al.*, 2018a) |
| Δ*flpA* | Δ*flpA::hyg^R^, pyrG*^+^ | This study |
| Δ*stk47* | Δstk47*::hyg^R^, pyrG*^+^ | This study |
| Δ*flpB* | Δ*flpB::hyg^R^, pyrG*^+^ | This study |
