## Supplementary material for "Identification and functional characterization of the putative members of the CTDK-1 kinase complex as regulators of growth and development in the genus *Aspergillus*": Table S4

**Table S4. Oligonucleotides used in this study.**

| **Primer** | **Sequence 5´-3´** | **Aim** |
| --- | --- | --- |
| **Oligonucleotides for *Aspergillus nidulans*** | | |
| PmtC-seq1 | CAGGAACACCTCGATCC | Sequencing of the  ORF of *An1459/pmtC* |
| PmtC-Up | GTGGGCTATGACGGC |  |
| PmtC-seq4 | CACCGGCGAACACAGC |  |
| PmtC-GSP2 | TTTCGCGAAGTGCAAGTCATAGCCAAGC |  |
| FlpA-PP1 | CAGCAGGCCACAACCATAGAAATACC | 3´-tagging and deletion of *An10640/flpA*  Constitutive expression of *An10640/flpA* |
| FlpA-PP2 | CATTCTGGAGCCGAGCTTGAAAATTTC |  |
| FlpA-GSP1 | CGAGACTCGAGGACTGCTCCCG |  |
| FlpA-GSP2 | TTCCTCGACCTCGTACTCTTCCATCTCAACC |  |
| FlpA-GSP3 | TGAAGCGTGGCGCTGACATACTCC |  |
| FlpA-GSP4 | GAAGGAGCCCAAAAGTCCATCTCAGC |  |
| FlpA-SMP1 | GAAATTTTCAAGCTCGGCTCCAGAATGACCGG  TCGCCTCAAACAATGCTCT |  |
| FlpA-GFP2 | GGAGTATGTCAGCGCCACGCTTCAGTCTGAGA  GGAGGCACTGATGCG |  |
| FlpA-GFP1 | GGTTGAGATGGAAGAGTACGAGGTCGAGGAA  GGAGCTGGTGCAGGCGCTGGAGCC |  |
| FlpA-seq1 | GATGTTCCAGATAGCAATCCCGC | Sequencing of the ORF of *An10640/flpA* |
| FlpA-seq2 | CGTGAGATTCTATGCGCGGCG |  |
| FlpA-seq3 | CTCCCGCGGTACACCAACTCG |  |
| FlpA-sPP1 | CGACTGGCTAGCCTTGAACATGACC | Diagnostic PCR reactions of the recombination at the *An10640/flpA locus* |
| FlpA-sGSP4 | CCGCTTTGTCGTCTGTACCTTAGAGC |  |
| FlpA-PP2´-ATG | GGAGCCGAGCTTGAAAATTTCCTCC | Constitutive expression of *An10640/flpA* |
| FlpA-gpdA-Up | GGAGGAAATTTTCAAGCTCGGCTCCCCATCCG  GTGCTCTGCACTCGACC |  |
| FlpA-gpdA-Up | CGAGTCTCGCTGCTGTTCAGGAGCCATCCATG  GTGATGTCTGCTCAAGCGGGG |  |
| FlpA-geneSP | ATGGCTCCTGAACAGCAGCGAGACTCG |  |
| Stk47-PP1 | GCTCCCACCACGGATCCTGG | 3´-tagging and deletion of *An8190/stk47*  Diagnostic PCR reactions of the recombination at the *An8190/stk47 locus* |
| Stk47-PP2 | CATTAATGAGGTGAGGGTGGCGG |  |
| Stk47-GSP3 | TAGGTGATTGACTATACCTCAAGTTCAGACAT  CTTTTGC |  |
| Stk47-GSP4 | GCCATGGGAGCTGTCGGG |  |
| Stk47-GSP1 | CGAATGCTAGCACGAGAACCGCC |  |
| Stk47-GSP2 | CTGCTGCCCAGGCTGCGGTACG |  |
| Stk47-sPP1 | GATAACGGGGAGGTGACTGGGG |  |
| Stk47-sGSP4 | CGAACGAGGCGAACGCACC |  |
| Stk47-SMP1 | CCGCCACCCTCACCTCATTAATGACCGGTCGC  CTCAAACAATGCTCT |  |
| Stk47-GFP1 | CGTACCGCAGCCTGGGCAGCAGGGAGCTGGTG  CAGGCGCTGGAGCC |  |
| Stk47-GFP2 | GCAAAAGATGTCTGAACTTGAGGTATAGTCAA  TCACCTAGTCTGAGAGGAGGCACTGATGCG |  |
| FlpB-PP1 | CCGAATGCAGTGCAGGAACTCC | 3´-tagging and deletion of *An6312/flpB*  Diagnostic PCR reactions of the recombination at the *An6312/flpB locus* |
| FlpB-PP2 | CATCGTGTCGAATTCTGCGCTTTTGC |  |
| FlpB-GSP3 | TAATTTGCATCTGGGCATATAGGGATTGACC |  |
| FlpB-GSP4 | GCAAATCCCAATAATCCCCAATCCG |  |
| FlpB-GSP1 | GATGGCAGATCCGTTTGAAGTTCGC |  |
| FlpB-GSP2 | AACAGCGTTGATCATCTCATTCCGC |  |
| FlpB-sPP1 | GTGGGCGGTCTCTATGGAGCG |  |
| FlpB-sGSP4 | GCGGATGAGAGAAGGAGCTCCAGC |  |
| FlpB-SMP1 | GCAAAAGCGCAGAATTCGACACGATGACCGG  TCGCCTCAAACAATGCTCT |  |
| FlpB-GFP1 | GCGGAATGAGATGATCAACGCTGTTGGAGCT  GGTGCAGGCGCTGGAGCC |  |
| FlpB-GFP2 | GGTCAATCCCTATATGCCCAGATGCAAATTAG  TCTGAGAGGAGGCACTGATGCG |  |
| **Oligonucleotides for *Aspergillus fumigatus*** | | |
| Fw_Cass_stk47 | GCCCACTGCCTTTATTTCATACTATATCTAGTTTTTAAAAAGCTTGCATGCCTGCAGGTC | Deletion of *stk47* |
| Rv_Cass_stk47 | GACAGGTATGGAGCATATATACGGCGCAGAAGTGCACAAACCGAGCTCCCAAATCTGTCC |  |
| Fw_Scr_stk47 | CGAATTGACTAGACGAAGGACA | Diagnostic PCR of deletion of *stk47* |
| Rv_Scr_stk47 | TGCTGTCATCTACCAGGAAAAG |  |
| gRNA5'_stk47 | AAAATTTCTTGTGGTGCATC | gRNAs for deletion of *stk47* |
| gRNA3'_stk47 | GCATGAATTTACTGTTCAAC |  |
| Fw_Cass_flpA | GAACAAGACGTGATCCGTCTCACGTGATATCCCGCCAATCAGCTTGCATGCCTGCAGGTC | Deletion of *flpA* |
| Rv_Cass_flpA | GTAGCGGATATGTCGCCTGTGTAGGGTGGTCTCGTGATTTCCGAGCTCCCAAATCTGTCC |  |
| Fw_Scr_flpA | CAATACTGCCCAGAGTGAGC | Diagnostic PCR of deletion of *flpA* |
| Rv_Scr_flpA | CTCTTGACTGAGCGAGCTGT |  |
| gRNA5'_flpA | CCAGCACCACAGGATGACTT | gRNAs for deletion of *flpA* |
| gRNA3'_flpA | AATGAATTGTCTTCTGGGAC |  |
| Fw_Cass_flpB | CTTTTTGAGGCAATCGGACTAGTTGAGCATTCTGAAAACTAGCTTGCATGCCTGCAGGTC | Deletion of *flpB* |
| Rv_Cass_flpB | ACACCGCCTATGAATATAGCATCAAGCCCAGGATATATAGCCGAGCTCCCAAATCTGTCc |  |
| Fw_Scr_flpB | CAGCTACCATCTTATCGCAGAC | Diagnostic PCR of deletion of *flpB* |
| Rv_Scr_flpB | GGGTTTCCACTCACTGACATTA |  |
| gRNA5'_flpB | TGCCATCATGGTTGCGGTTC | gRNAs for deletion of *flpB* |
| gRNA3'_flpB | ATTGGTGTTAGGGAGTTTTG |  |
